# An Autoantigen Profile of Human A549 Lung Cells Reveals Viral and Host Etiologic Molecular Attributes of Autoimmunity in COVID-19

**DOI:** 10.1101/2021.02.21.432171

**Authors:** Julia Y. Wang, Wei Zhang, Michael W. Roehrl, Victor B. Roehrl, Michael H. Roehrl

## Abstract

We aim to establish a comprehensive COVID-19 autoantigen atlas in order to understand autoimmune diseases caused by SARS-CoV-2 infection. Based on the unique affinity between dermatan sulfate and autoantigens, we identified 348 proteins from human lung A549 cells, of which 198 are known targets of autoantibodies. Comparison with current COVID data identified 291 proteins that are altered at protein or transcript level in SARS-CoV-2 infection, with 191 being known autoantigens. These known and putative autoantigens are significantly associated with viral replication and trafficking processes, including gene expression, ribonucleoprotein biogenesis, mRNA metabolism, translation, vesicle and vesicle-mediated transport, and apoptosis. They are also associated with cytoskeleton, platelet degranulation, IL-12 signaling, and smooth muscle contraction. Host proteins that interact with and that are perturbed by viral proteins are a major source of autoantigens. Orf3 induces the largest number of protein alterations, Orf9 affects the mitochondrial ribosome, and they and E, M, N, and Nsp proteins affect protein localization to membrane, immune responses, and apoptosis. Phosphorylation and ubiquitination alterations by viral infection define major molecular changes in autoantigen origination. This study provides a large list of autoantigens as well as new targets for future investigation, e.g., UBA1, UCHL1, USP7, CDK11A, PRKDC, PLD3, PSAT1, RAB1A, SLC2A1, platelet activating factor acetylhydrolase, and mitochondrial ribosomal proteins. This study illustrates how viral infection can modify host cellular proteins extensively, yield diverse autoantigens, and trigger a myriad of autoimmune sequelae.

## Introduction

To gain better understanding of the transient and chronic autoimmune symptoms caused by SARS-CoV-2 infection, we have embarked on an endeavor to establish a comprehensive autoantigenome for COVID-19. In a previous study, we identified a repertoire of autoantigens (autoAgs) from human fetal lung fibroblast HFL1 cells that are strongly tied to neurological and diverse autoimmune symptoms of COVID- 19 (1). In this study, we aim to identify additional autoAgs from human lung epithelium-like A549 cells, an adenocarcinoma cell line that is frequently used as a model host in SARS-CoV-2 infection studies.

AutoAgs were identified based on the unique affinity between autoAgs and the glycosaminoglycan dermatan sulfate (DS) that we have discovered (2, 3). AutoAgs and DS form affinity complexes that can engage strong dual BCR signaling in autoreactive B1 cells to induce autoantibody production (4). Hence, any self-molecule capable of forming affinity complexes with DS has a high propensity to become autoantigenic. This unifying mechanism of autoantigenicity explains how seemingly unrelated self-molecules can all induce autoimmune B cell responses via a similar immunological signaling event. Based on DS-autoAg affinity, we have cataloged several hundred autoAgs from various cells and tissues (1, 5-7).

COVID-19 is accompanied by a wide range of autoimmune symptoms, including multisystem inflammatory syndrome in children, immune thrombocytopenic purpura, antiphospholipid syndrome, autoimmune cytopenia, immune-mediated neurological syndromes, Guillain-Barré syndrome, connective tissue disease-associated interstitial lung disease, autoimmune hemolytic anemia, autoimmune encephalitis, systemic lupus erythematosus, optic neuritis and myelitis, and acquired hemophilia (8-15). Numerous autoantibodies have been identified in COVID patients, including the classical ANA (antinuclear antibody) and ENA (extractable nuclear antigen) that are hallmarks of systemic autoimmune diseases, as well as others such as anti-neutrophil cytoplasmic antibody, lupus anticoagulant, antiphospholipid, anti-IFN, anti-myelin oligodendrocyte glycoprotein, and anti-heparin-PF4 complex antibodies (8-15).

SARS-CoV-2, or viruses in general, are opportunistic intracellular pathogens that rely on the host for replication and survival. They hijack the host transcription and translation machinery for their replication, they compromise the host immune defense to evade destruction, and they modulate the host cell cycle and apoptosis for symbiosis. These viral processes are accomplished through extensive modification of host cellular components, which also results in changes in self-molecules and the emergence of autoAgs. In our previous studies, we reported that self-molecules derived from apoptotic cells display strong affinity to DS, becoming a major source of autoAgs (2, 3). In this study, we report several important molecular mechanisms in SARS-CoV-2 infection that change host self-molecules to autoAgs, including direct interaction with viral components, perturbation by viral protein expression, and post-translational protein modification by ubiquitination and phosphorylation from viral infection.

## Results and Discussion

### A putative A549 autoantigenome identified by DS-affinity

By DS-affinity fractionation and mass spectrometry sequencing, we identified a global putative autoantigenome of 348 proteins from A549 cellular protein extracts, with 214 protein having strong affinity and 134 having intermediate affinity (Table 1). To find out whether these DS-affinity proteins are known autoAgs, we conducted an extensive literature search and confirmed that 198 (56.0%) proteins are known humoral autoAgs, with their specific autoantibodies reported in a wide spectrum of autoimmune diseases and cancers (see autoAg confirmatory references in Table 1). The remaining 150 proteins may be yet-to-be discovered putative autoAgs and await further investigation. For example, many ribosomal proteins are known autoAgs, but the 24 mitochondrial ribosomal proteins we identified have not yet been reported as autoAgs; given their structural similarity to ribosomal protein autoAgs, it is highly likely that mitochondrial proteins are a group of undiscovered autoAgs.

**Table 1.**
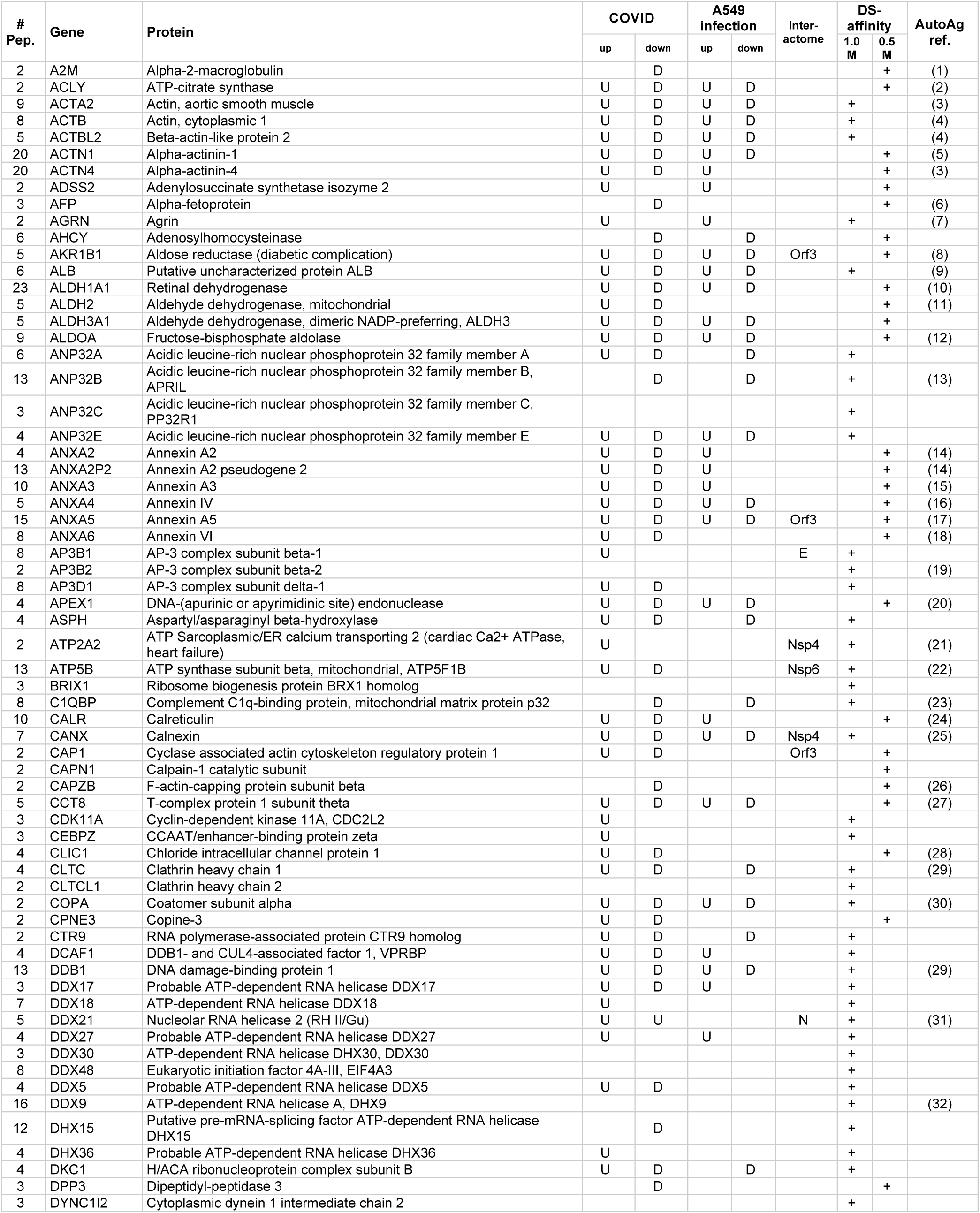
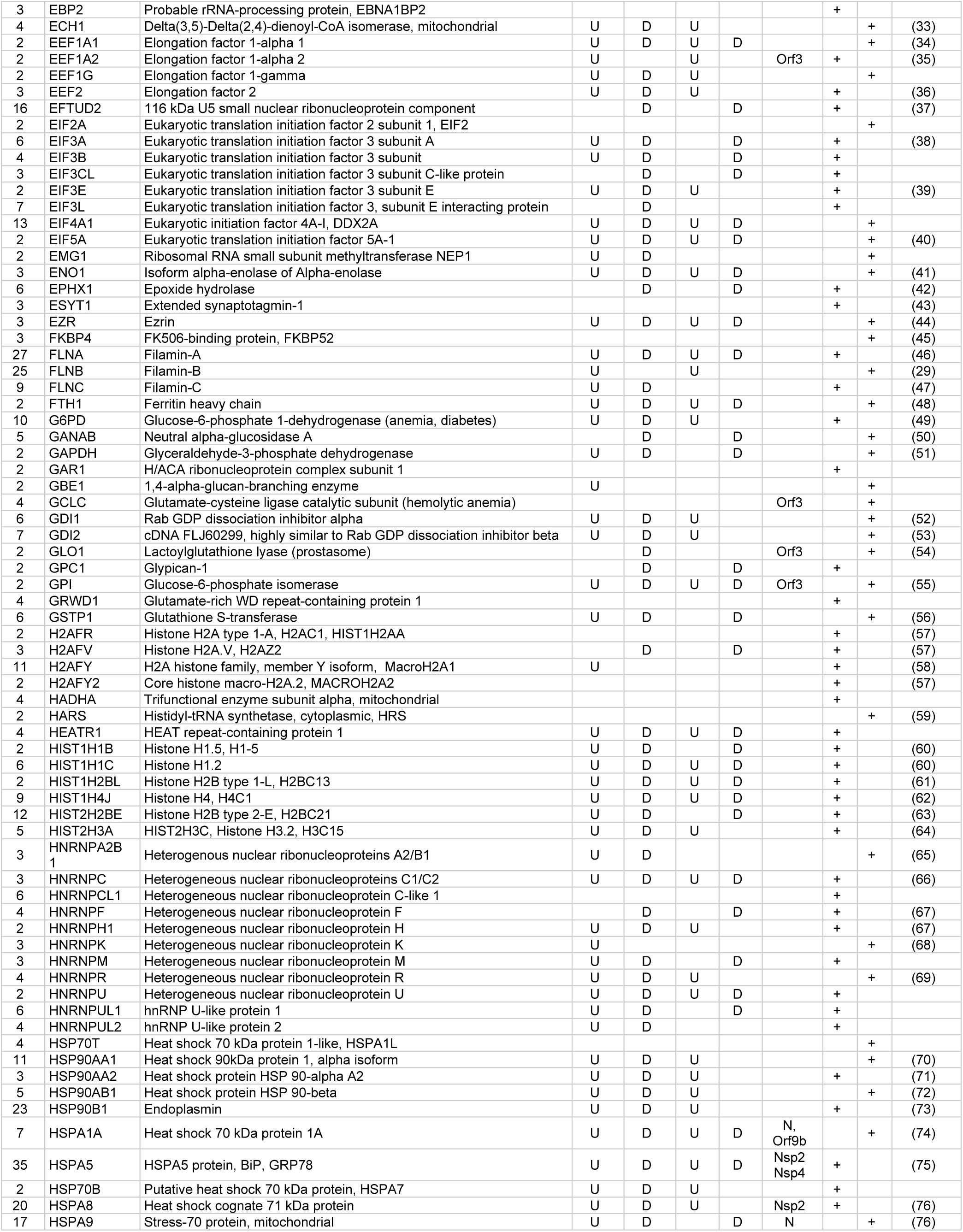
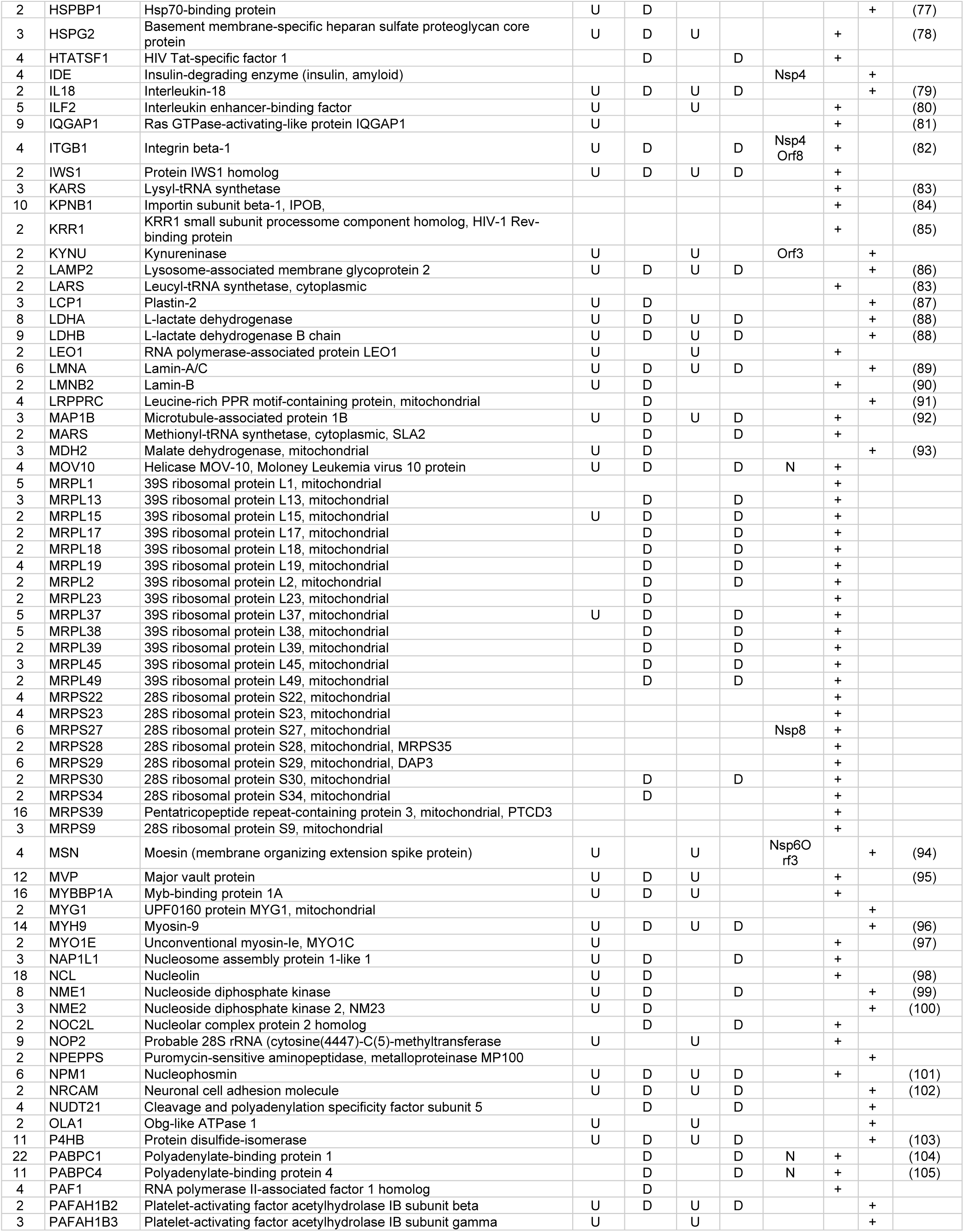
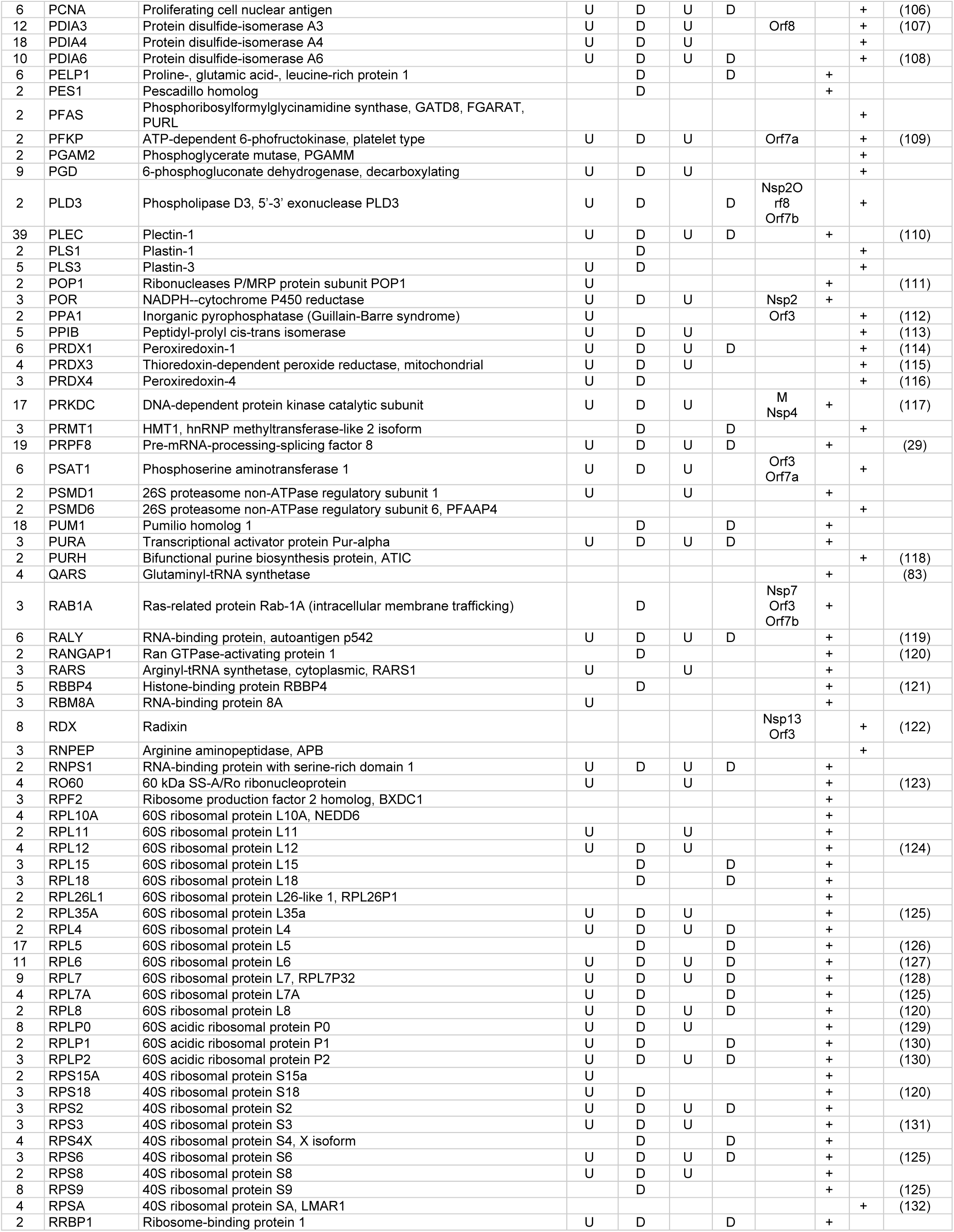
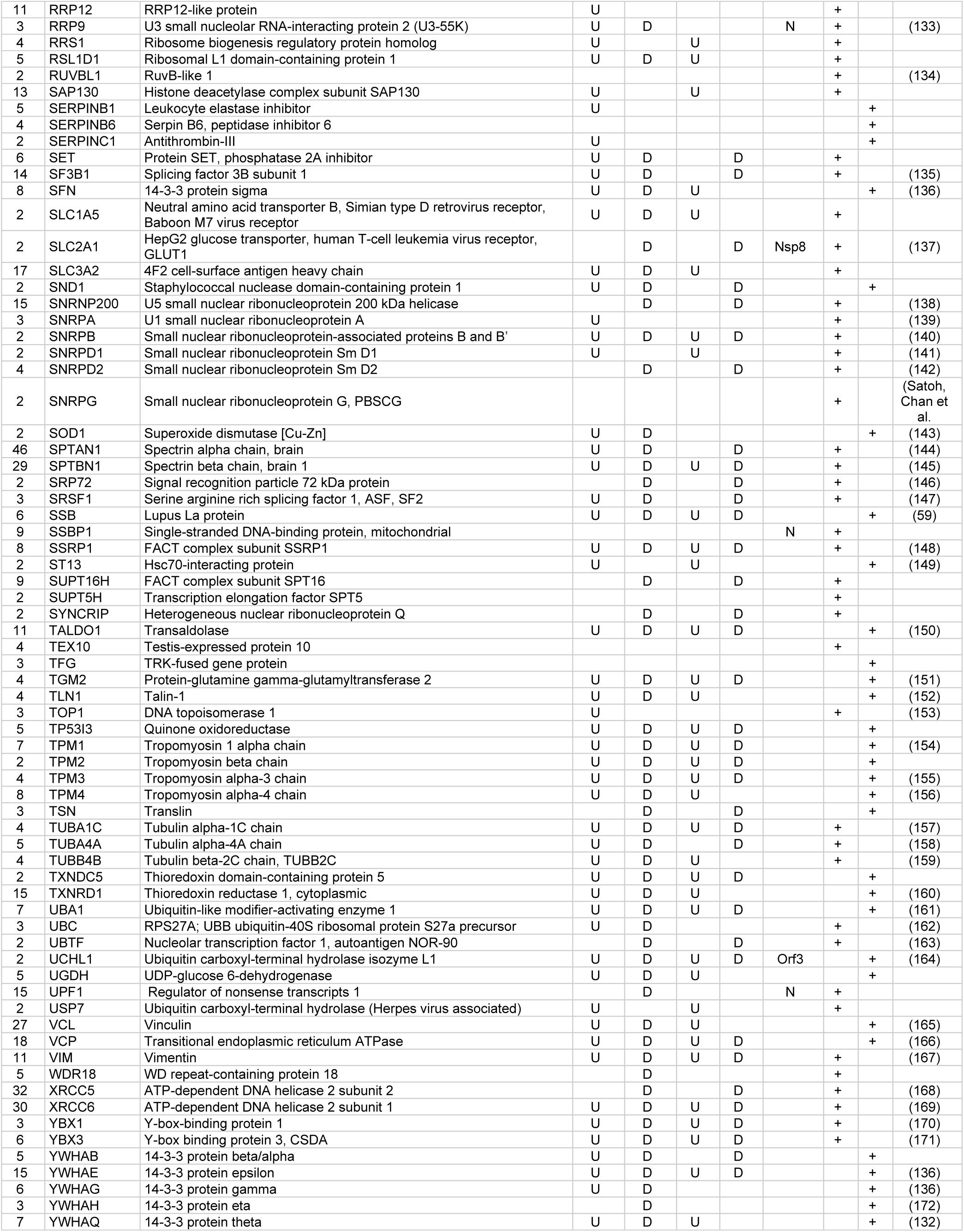
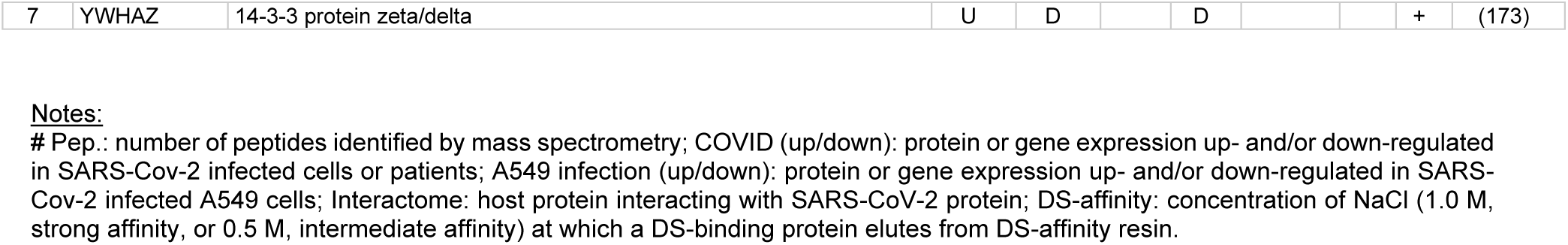
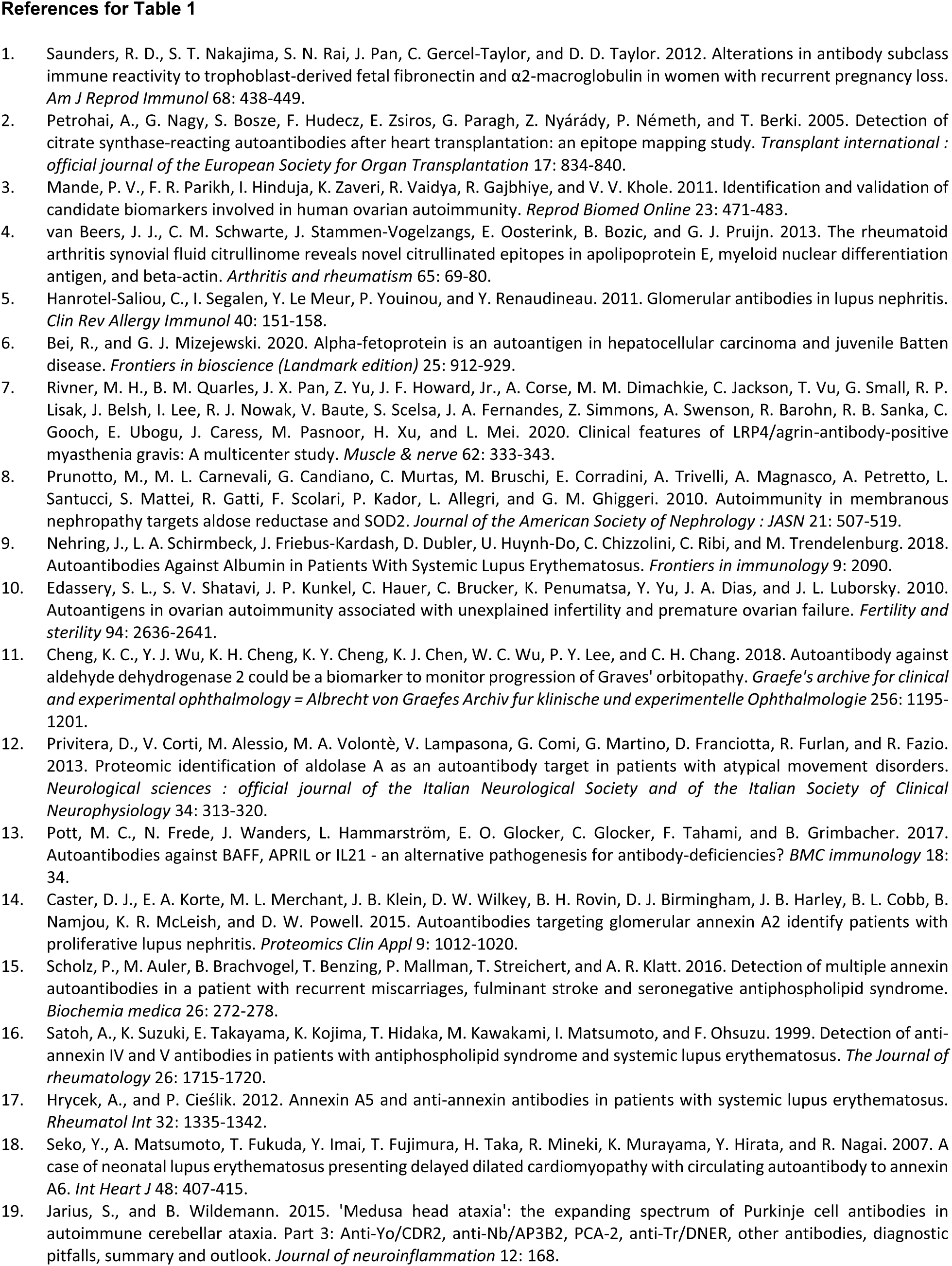
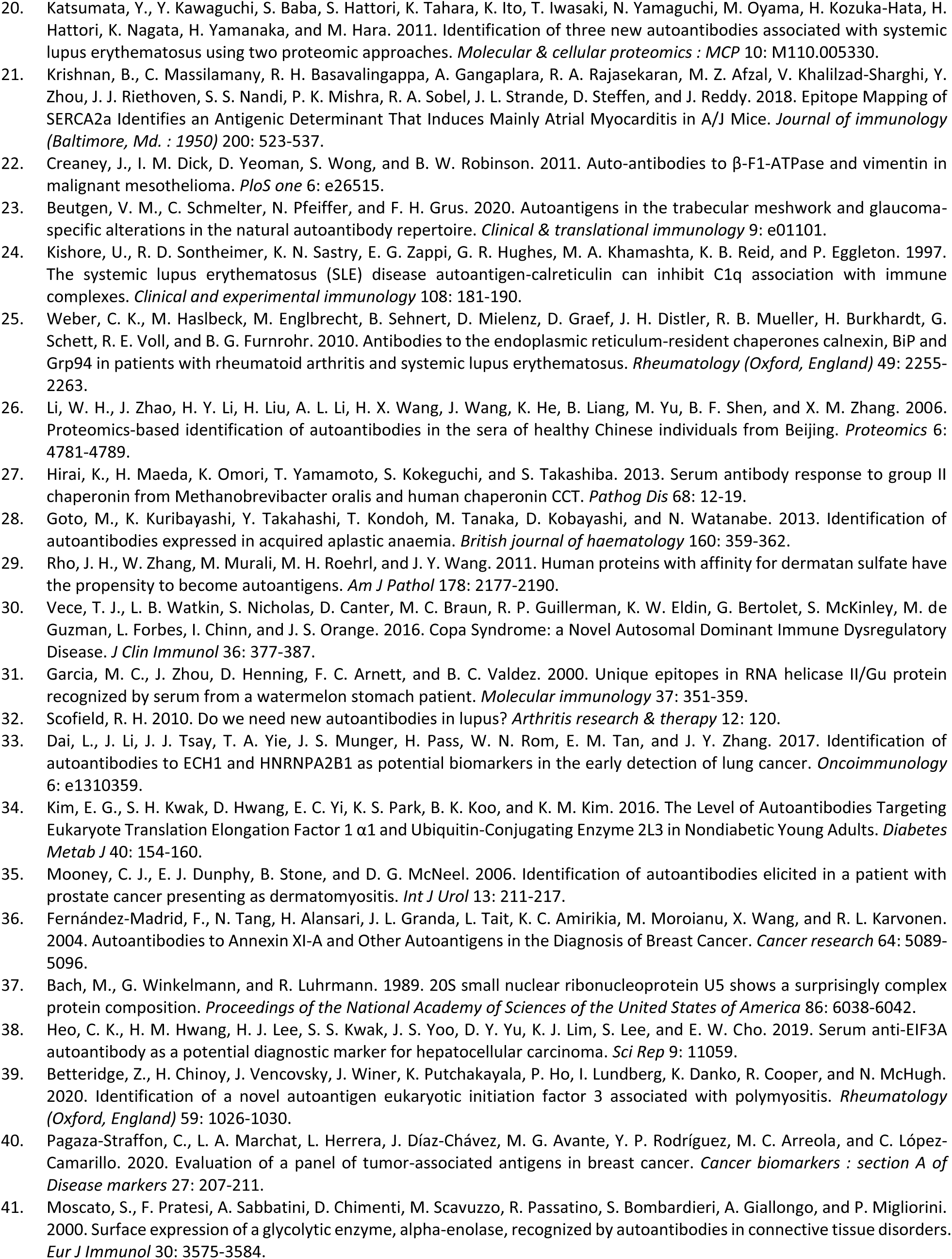
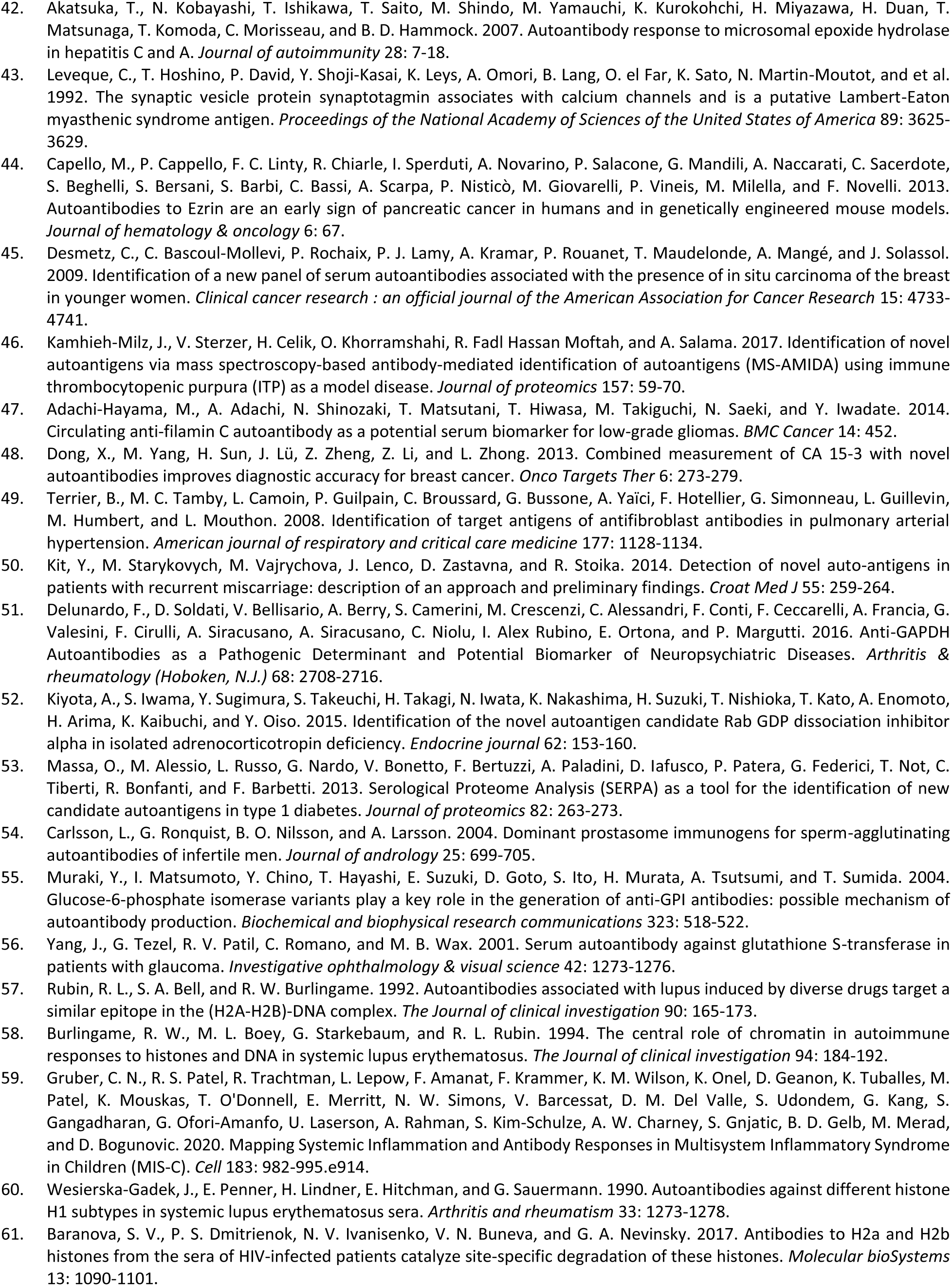
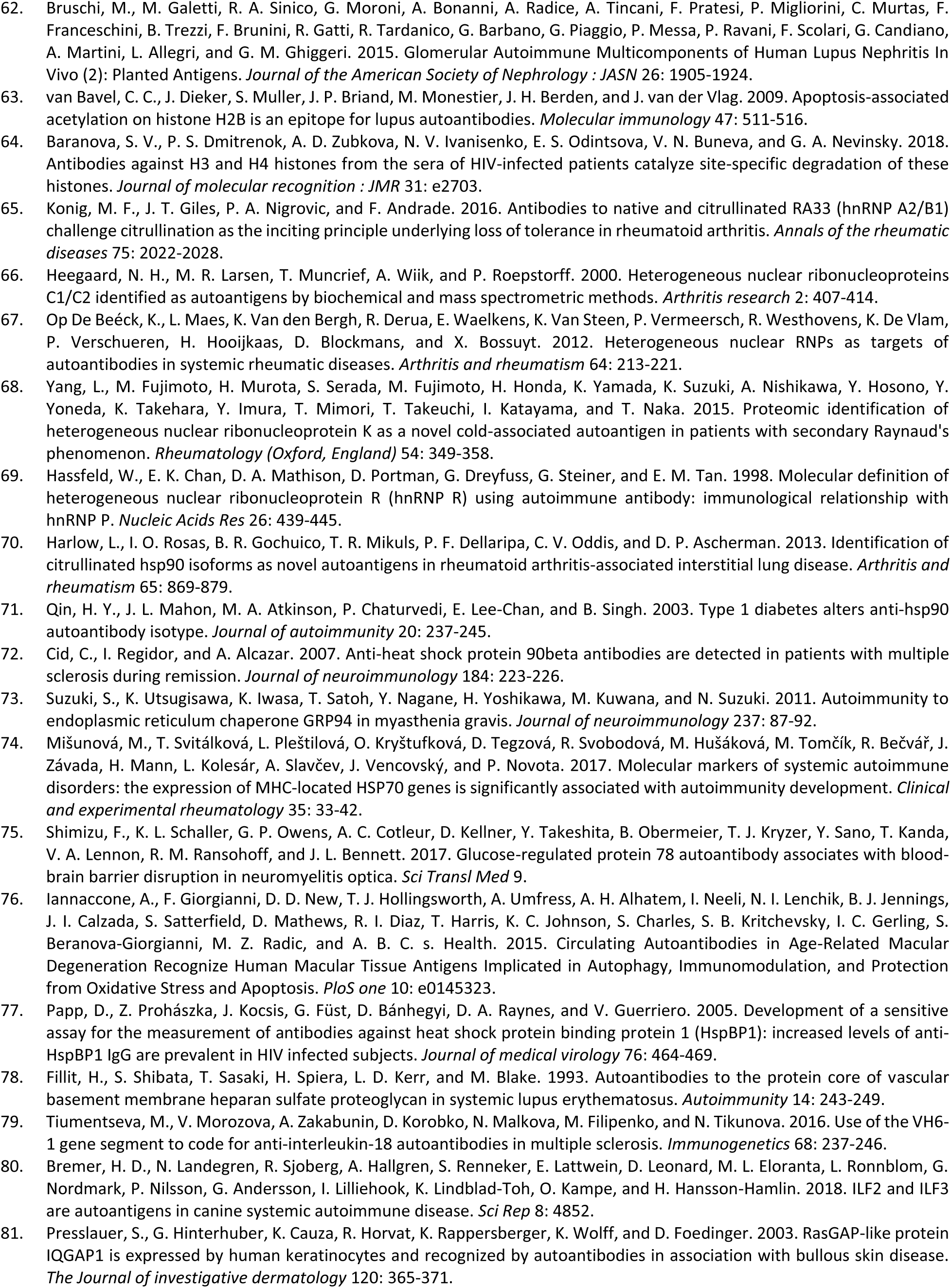
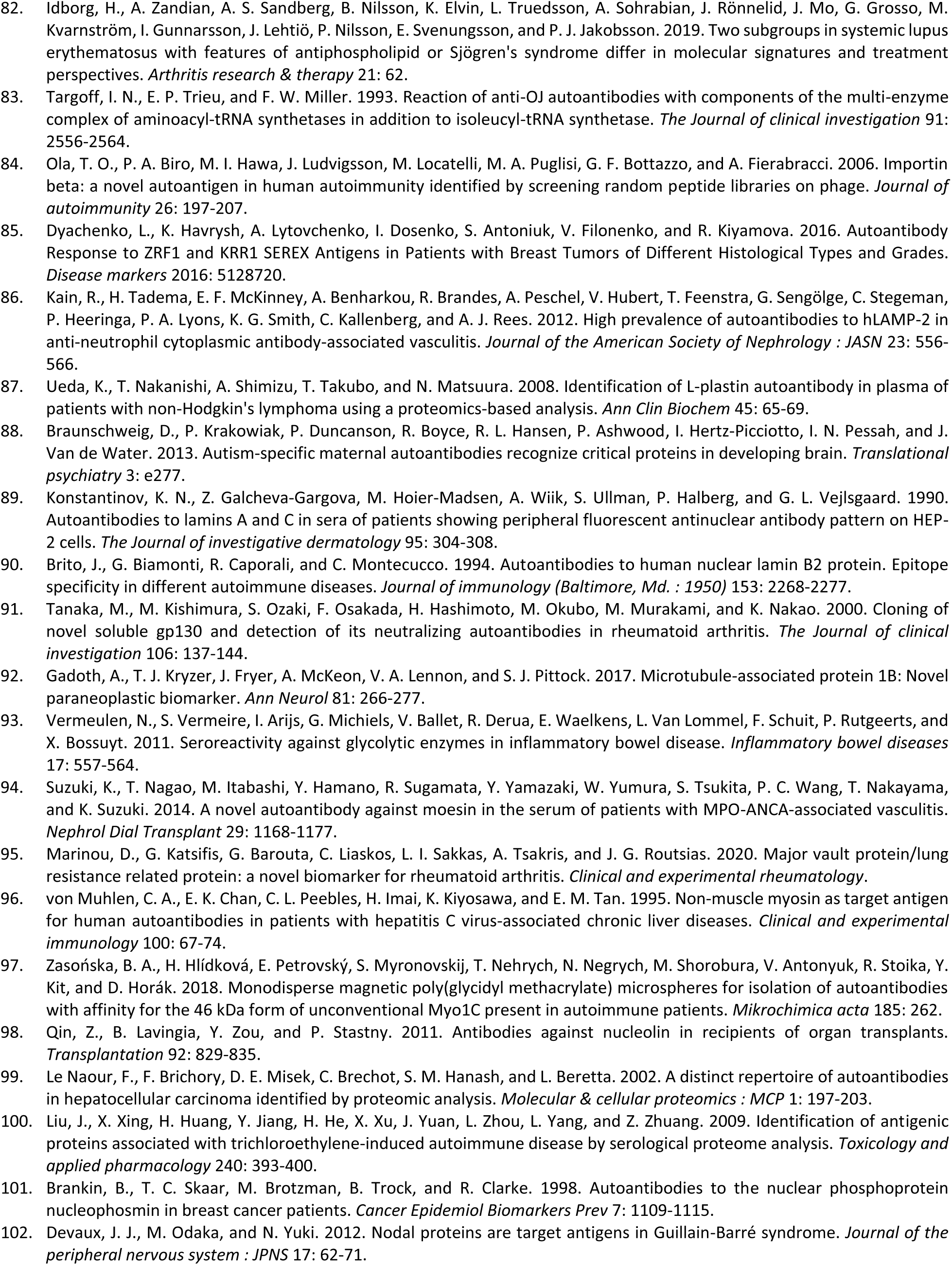
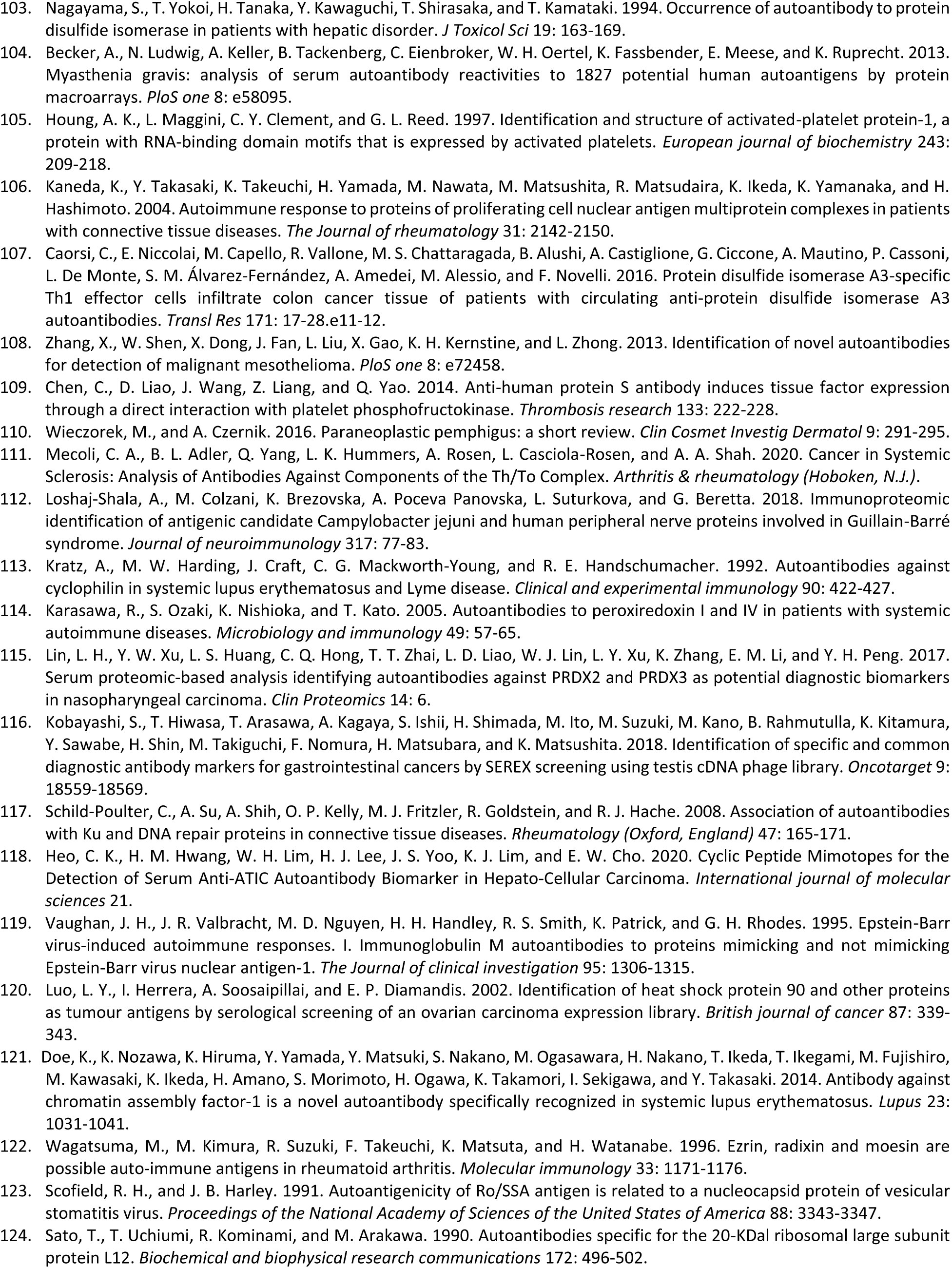
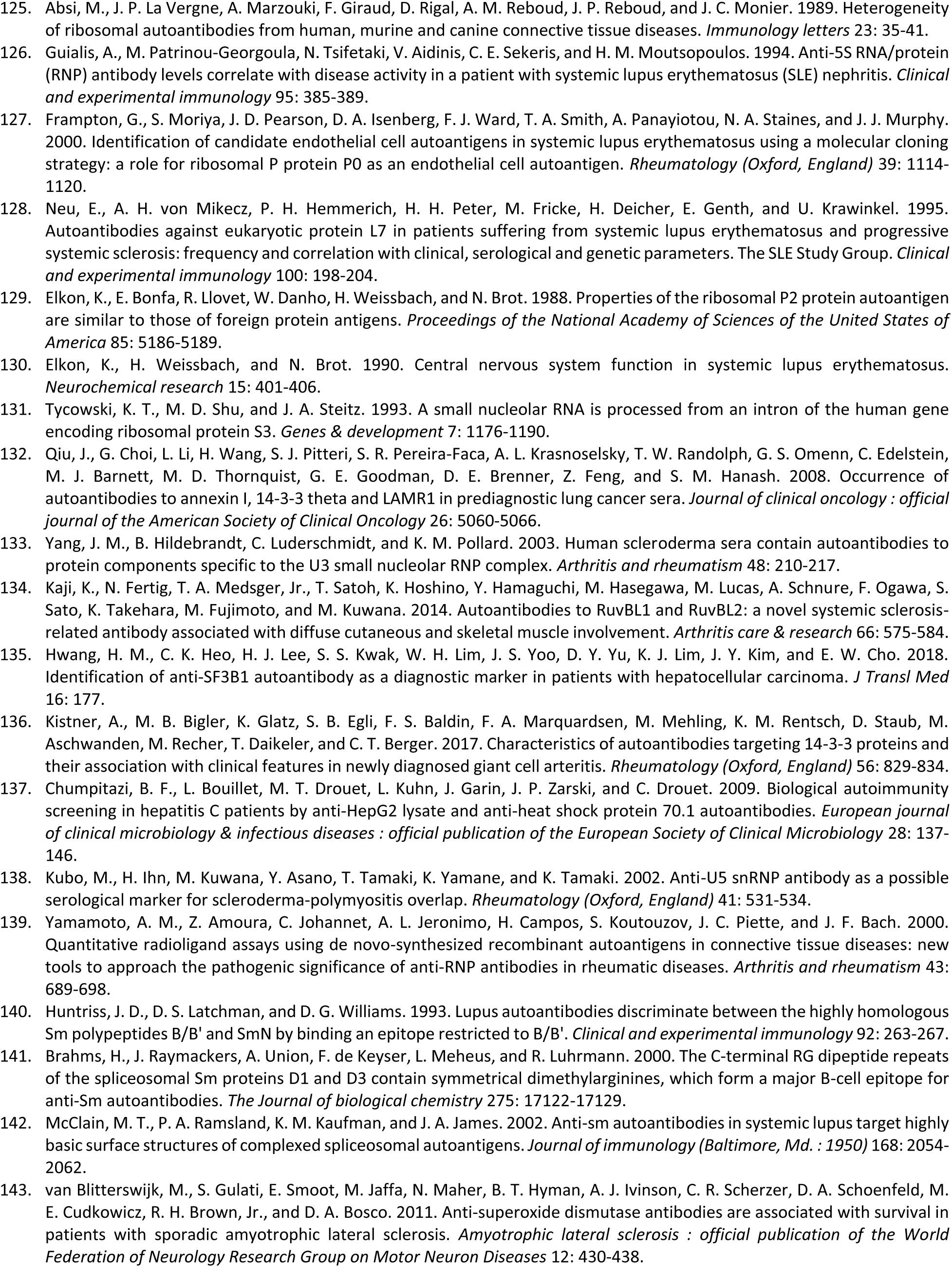
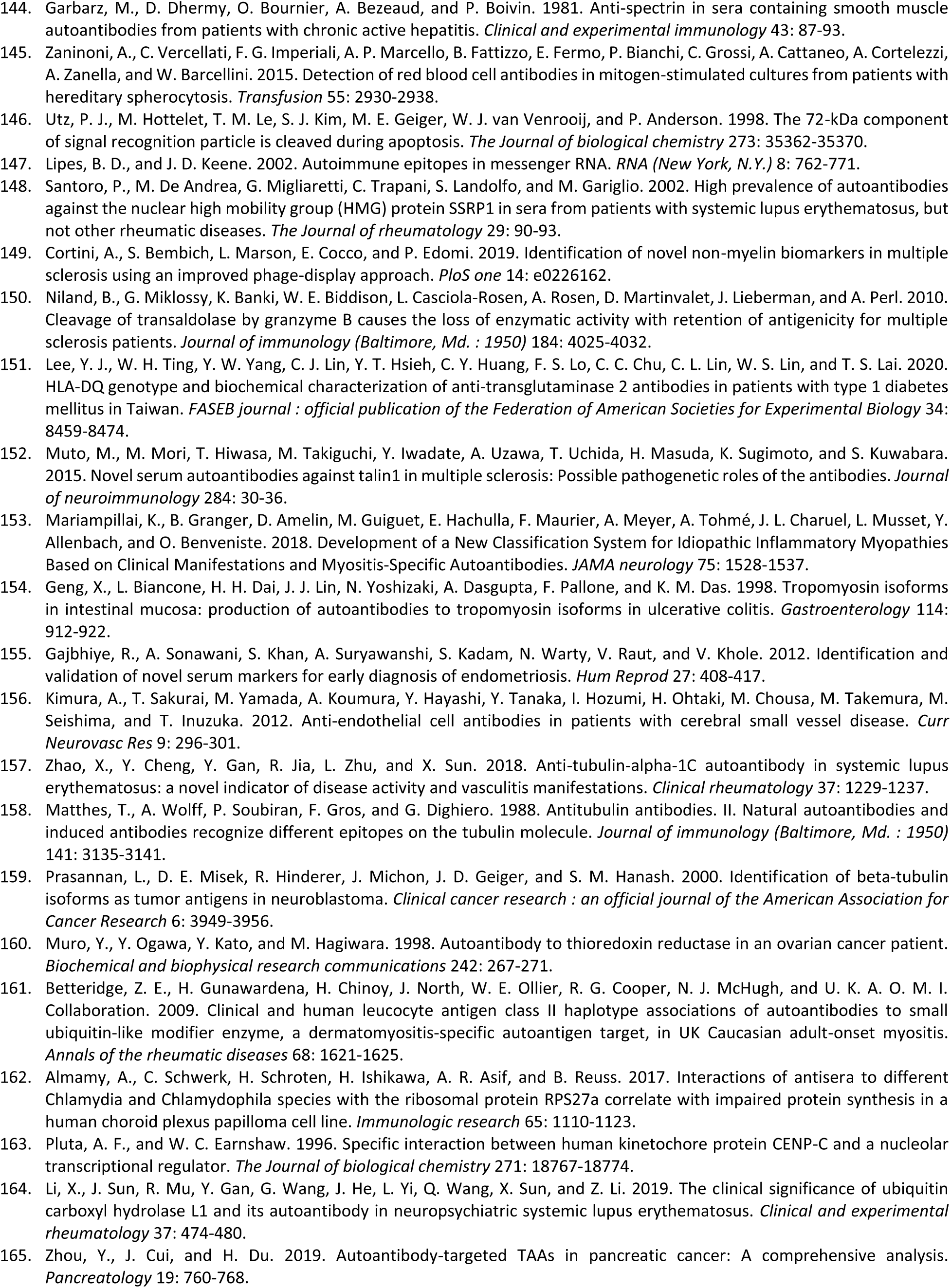
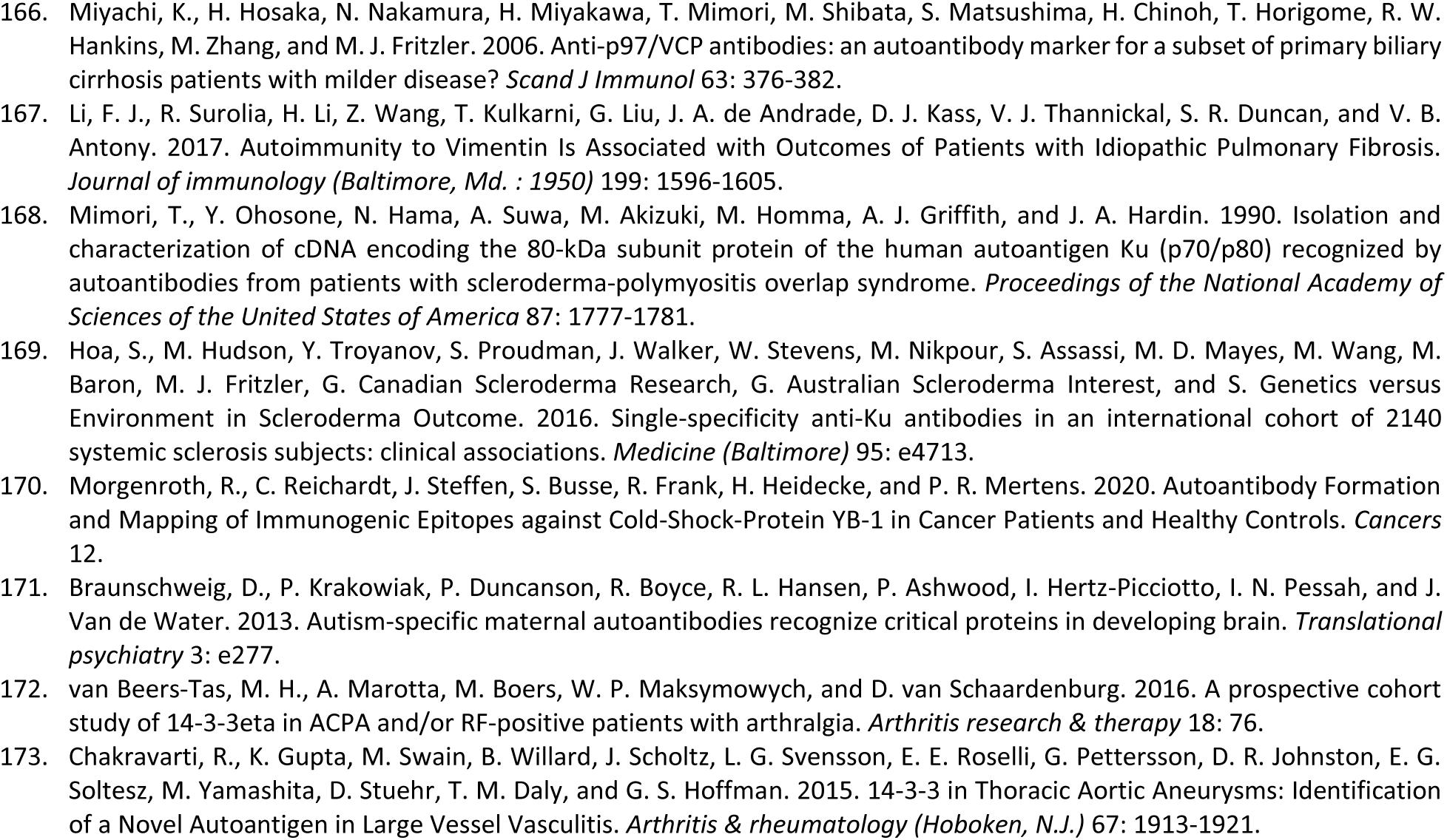
DS-affinity autoantigenome from human A549 cells

The 348 DS-affinity proteins are highly connected (Fig. 1). They exhibit 6,271 interactions, whereas a random set of 348 proteins is expected to have 2,536 interactions, as revealed by protein-protein interaction STRING analysis (16). The tight connections suggest that these known and putative autoAgs are originating from common biological pathways or processes. Our analysis shows that they are indeed predominantly associated with translation, mRNA metabolic process, ribonucleoprotein complex biogenesis, vesicle and vesicle-mediated transport, chromosome, and cytoskeleton (Fig. 1).

**Fig. 1.**
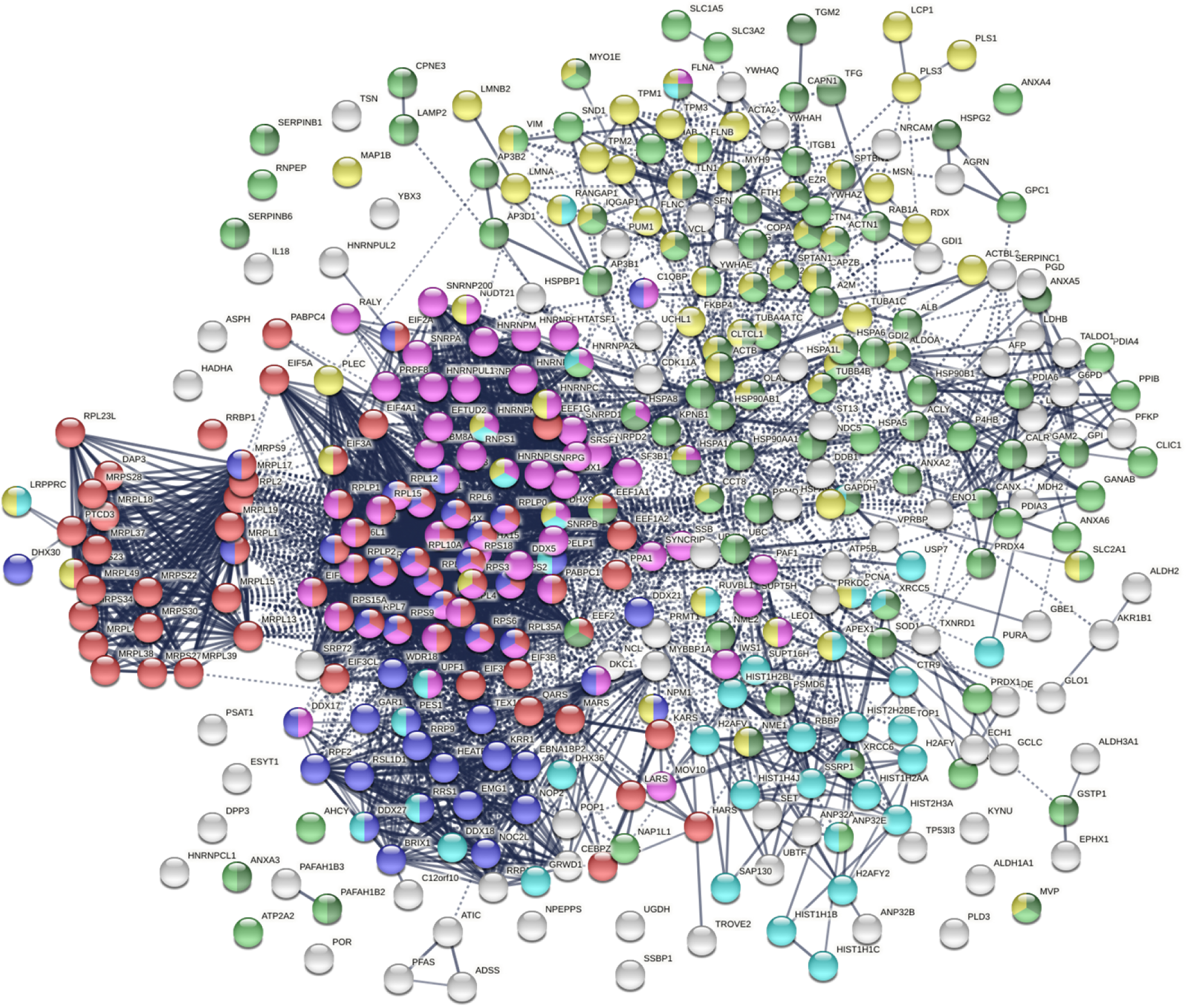
The autoantigenome from A549 cells identified by DS affinity. Marked proteins are associated with translation (69 proteins, red), mRNA metabolic processing (69 proteins, pink), ribosome biogenesis (43 proteins, blue), vesicle (87 proteins, green) and vesicle-mediated transport (72 proteins, dark green), chromosomes (40 proteins, aqua), and cytoskeleton (65 proteins, gold).

### COVID-altered proteins among the A549 autoantigenome

To find out whether the known and putative autoAgs identified by DS-affinity may play a role in SARS-CoV-2 infection, we compared our A549 autoantigenome with currently available COVID data compiled in the Coronascape database (17-37). Of our 348 autoantigenome proteins from A549 cells, 291 (83.6%) have been found to be COVID-altered, i.e., up- and/or down-regulated at protein and/or mRNA level in SARS-CoV-2 infected cells or patient tissues (Table 1). Because the COVID data have been generated from various research labs using different techniques and sources of cells or tissues, 190 proteins are found to be up in some studies but down in others. In total, 231 proteins are found up-regulated, and 252 are found down-regulated in SARS-CoV-2 infection. Based on reported autoantibodies, 191 (65.6%) COVID-altered proteins are confirmed autoAgs (Table 1).

Based on gene ontology (GO) cellular component analysis, proteins of the A549 DS-affinity autoantigenome that are also altered in COVID infection can be located to membrane-bound organelles (247 proteins), nucleus (177 proteins), ribonucleoprotein complex (95 proteins), mitochondrion (46 proteins), endoplasmic reticulum (45 proteins), secretory granules (41 proteins), melanosome (27 proteins), myelin sheath (28 proteins), and axon (16 proteins).

Within the total A549 autoantigenome, the 291 COVID-altered proteins form a tightly interacting network (Fig. 2). At high STRING protein-protein interaction confidence level, these proteins exhibit 2,249 interactions, whereas 953 interactions would be expected of a random collection of proteins of the same size. By GO biological process analysis, the COVID-altered proteins are significantly enriched in various biological processes, including translation, peptide biosynthetic process, RNA catabolic process, nucleobase-containing compound catabolic process, SRP-dependent cotranslational protein targeting to membrane, protein localization to organelle, and symbiont process. Among these processes associated with COVID-altered proteins, the hierarchical cluster tree root points to ribonucleoprotein complex biogenesis (Fig. 3).

**Fig. 2.**
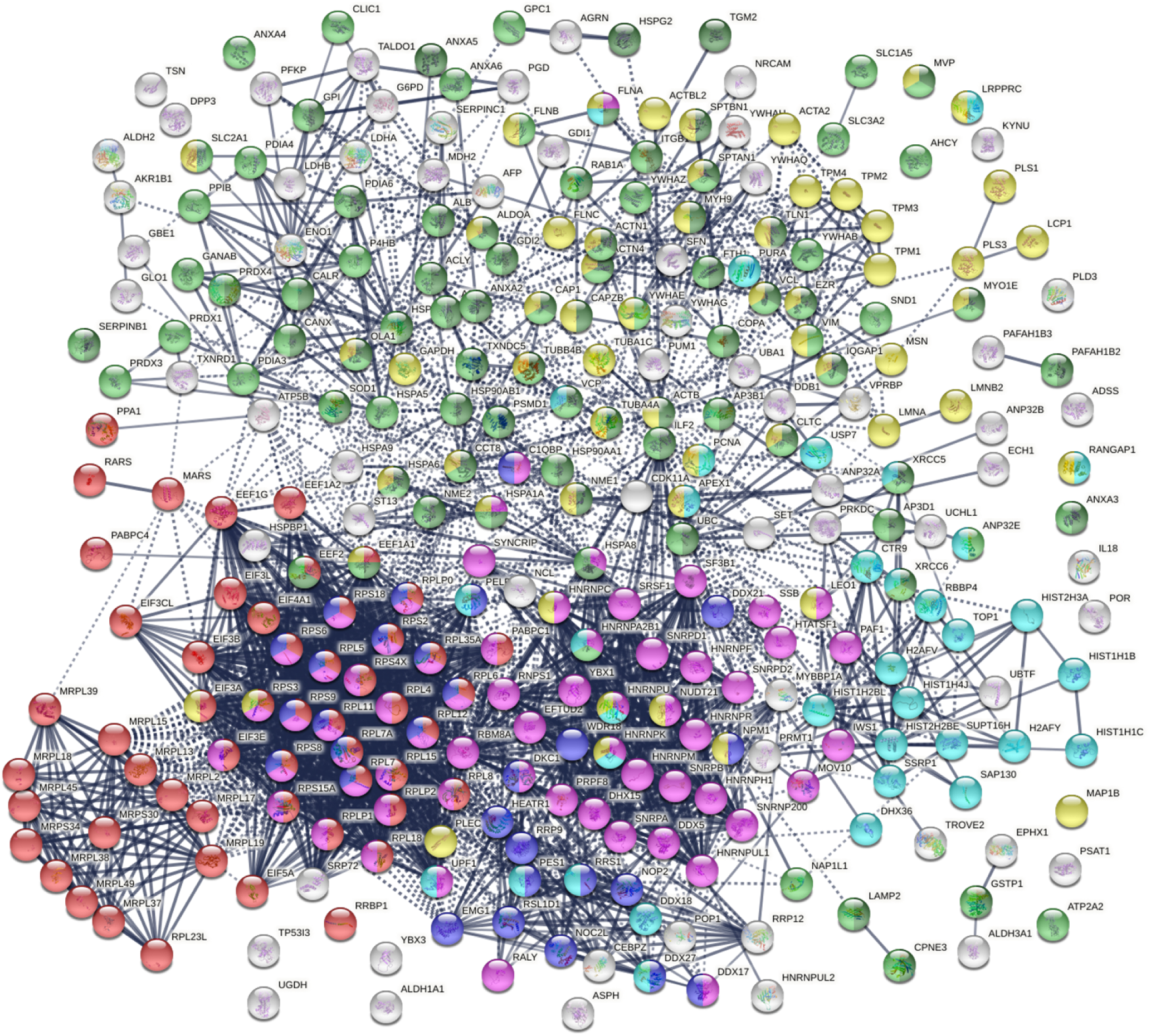
COVID-altered proteins shared with the A549 autoantigenome. Marked proteins are associated with translation (53 proteins, red), mRNA metabolic process (63 proteins, pink), vesicle (79 proteins, green) and vesicle-mediated transport (64 proteins, dark green), cytoskeleton (58 proteins, gold), chromosomes (35 proteins, aqua), and ribosome biogenesis (29 proteins, blue).

**Fig. 3.**
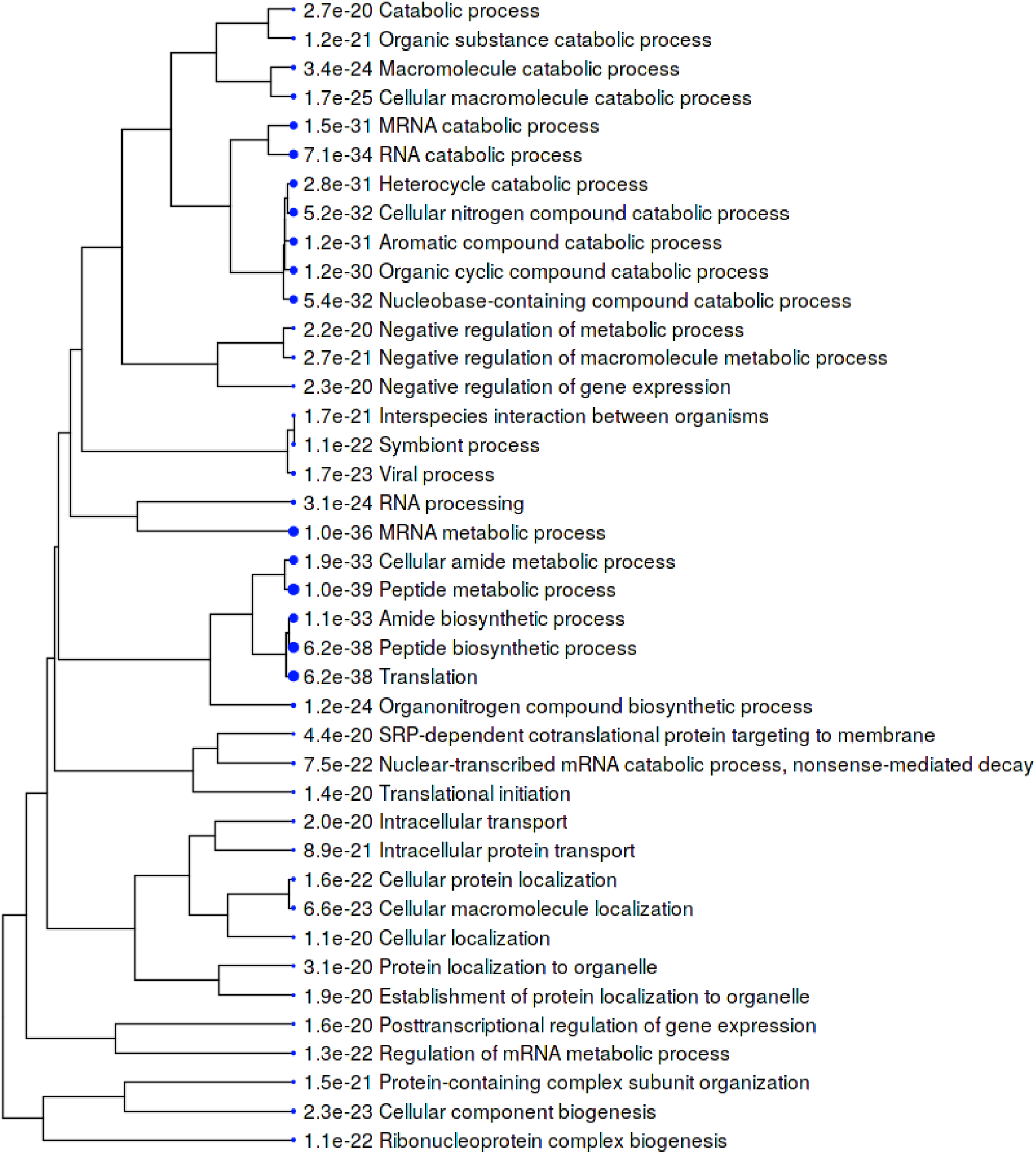
Top 40 enriched GO biological processes among COVID-altered proteins shared with the A549 autoantigenome. Bigger dots indicate more significant P-values.

Combined pathway and process enrichment analyses also show that the COVID-altered DS-affinity proteins are most significantly related to peptide biosynthetic process, metabolism of RNA, and ribonucleoprotein complex biogenesis (Fig. 4). The up-regulated autoAgs are more related to Nop56p- associated pre-rRNA complex, RNA catabolic process, and vesicle-mediated transport, whereas the down- regulated autoAgs are more related to translation and ribosome biogenesis. COVID-altered autoAgs are also significantly associated with regulated exocytosis, platelet degranulation, smooth muscle contraction, and IL-12 signaling.

**Fig. 4.**
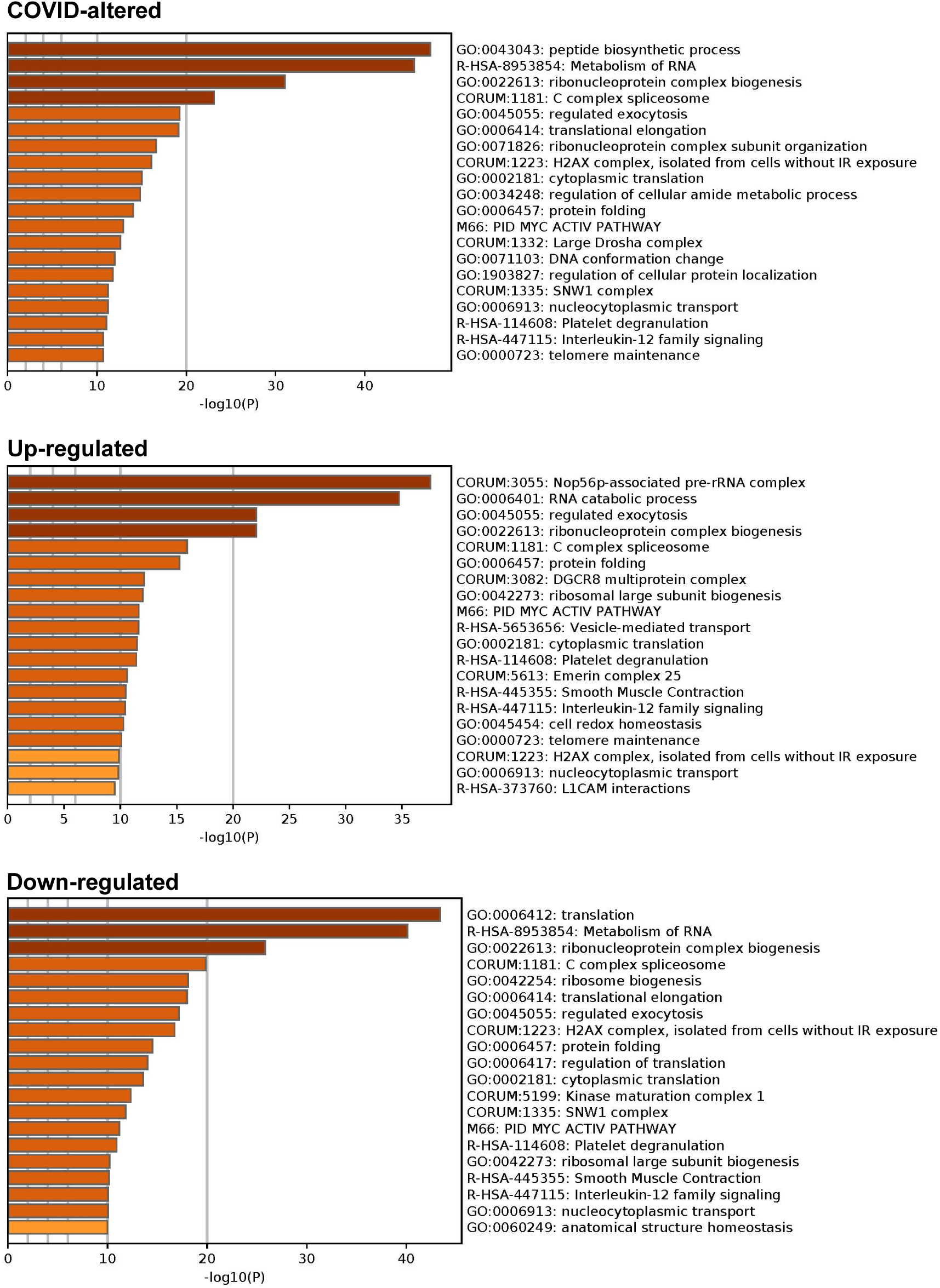
Top 20 enriched pathways and processes among COVID-altered autoAgs. Top: 298 COVID- altered autoAgs. Middle: 231 up-regulated autoAgs in COVID. Bottom: 252 down-regulated autoAgs in COVID.

The molecular functions of the COVID-altered DS-affinity autoAgs include RNA binding (76 proteins), hydrolase activity (58 proteins), purine ribonucleotide triphosphate binding (55 proteins), pyrophosphatase activity (35 proteins), nucleoside-triphosphatase activity (34 proteins), oxidoreductase activity (26 proteins), ATPase activity (25 proteins), ubiquitin protein ligase binding (21 proteins), mRNA 3’-UTR binding (11 proteins) and 5’-UTR binding (3 proteins), and helicase activity (13 proteins).

### AutoAgs that interact with SARS-CoV-2 proteins

To study the origins of COVID-induced autoAg alterations, we examined DS-affinity proteins that are involved in the interactomes of SARS-CoV-2 viral proteins (19, 30, 34) and found 38 DS-affinity proteins that interact directly with different viral proteins, with 25 of them being known autoAgs, e.g., ATP5B, CANX, DDX21, EEF1A2, PDIA3, and SLC2A1 (Table 1). Orf3 protein is the most striking, as its interactome includes 14 DS-affinity proteins and 9 known autoAgs. N protein interacts with 9 DS-affinity proteins and 6 known autoAgs, including 2 helicases (DDS21 and MOV10), 2 poly(A)-binding proteins (PABPC1 and PABPC4). Nsp4 interact with 6 DS-affinity proteins and 5 known autoAgs. Nsp4 interacts with IDE (insulin degrading enzyme), which is not a known autoAg.

A few host DS-affinity proteins interact with more than one SARS-CoV-2 protein. RAB1A (involved in intracellular membrane trafficking) interacts with 3 viral proteins (Orf3, Orf7b, and Nsp7), and PLD3 (phospholipase) also interacts with 3 viral proteins (Nsp2, Orf7b, and Orf8), but neither RAB1A nor PLD3 has been discovered as autoAgs. HSPA5 (GRP78/BiP) interacts with Nsp2 and Nsp4, HSPA1A interacts with N and Orf9b, and both heat shock proteins are known autoAg. PRKDC, a known autoAg, interacts with M and Nsp. Interestingly, of the ezrin-radixin-moesin protein family that connects the actin cytoskeleton to the plasma membrane, RDX is found to interact with Orf3 and Nsp13, MSN interacts with Orf3 and Nsp6, and EZR is COVID-altered. Furthermore, RDX, MSN, and EZR are all known autoAgs (Table 1).

### AutoAgs from perturbation by viral protein expression

To find out how individual SARS-CoV-2 proteins affect the host, Stukalov et al. conducted extensive proteomic analysis of A549 cells transduced to express individual SARS-CoV-2 proteins (34). By comparing with their data, we identified 167 DS-affinity proteins that are perturbated by viral protein expression in A549 cells. Among all SARS-CoV-2 proteins, Orf3 expression produced the largest number of potential autoAgs, with 26 up- and 36 down-regulated being DS-affinity proteins (Fig. 5, Supplemental Table 1). Other viral protein expressions affected various numbers of DS-affinity proteins, including E (20 up and 18 down), Orf9b (6 up and 16 down), M (10 up and 11 down), N (10 up and 8 down), Nsp13 (5 up and 11 down), and Nsp12 (3 up and 12 down). Interestingly, S expression yielded only 3 up- and 2 down-regulated DS-affinity proteins, suggesting that it may not have much intracellular activity.

**Fig. 5.**
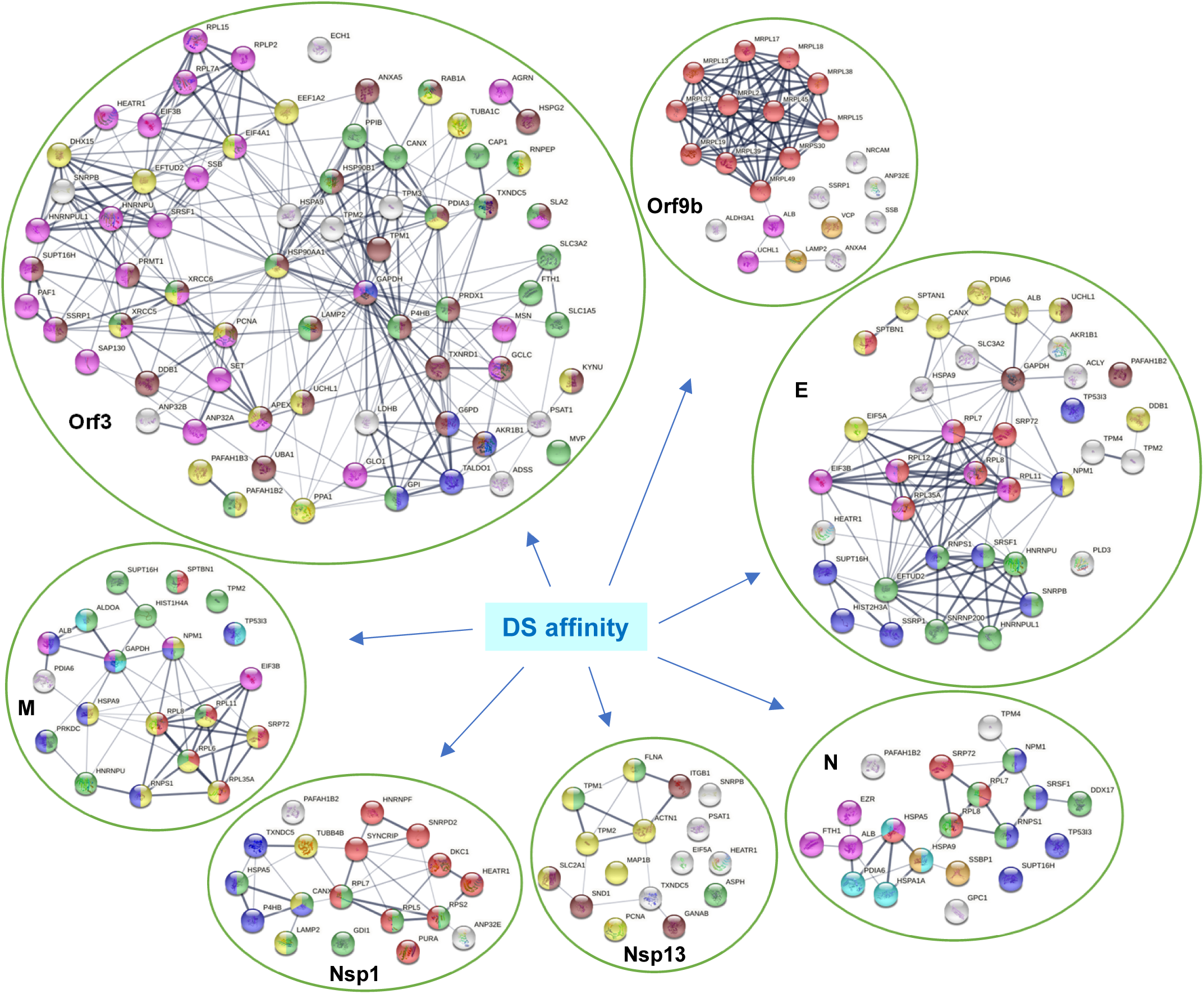
DS-affinity proteins that interact with SARS-CoV-2 viral proteins or are perturbed in A549 cells expressing individual viral proteins. **Orf3:** regulation of gene expression (pink), cytoplasmic vesicle (green), monosaccharide biosynthetic process (blue), response to stress (brown), and hydrolase activity (gold). **Orf9b:** mitochondrial translation (red), mitochondrion localization (pink), autophagosome maturation (gold). **E:** establishment of protein localization to membrane (red), translation initiation (pink), mRNA splicing (green), regulation of macroautophagy (brown), post-translational protein modification (gold), RNA polymerase II transcription (blue). **M:** establishment of protein localization to membrane (red), intracellular protein transport (gold), organelle organization (green), nicotinamide nucleotide metabolic process (aqua), regulation of apoptotic process (blue), and symbiont process (pink). **N:** maintenance of location in cell (pink), protein localization to ER (red), protein folding (aqua), mitochondrial nucleoid (amber), RNA polymerase II transcription (blue), and RNA-binding (green). **Nsp1:** protein localization (green), gene expression (red), protein processing in ER (blue), and phagosome (gold). **Nsp13:** cytoskeleton (gold), regulation of muscle contraction (green), and melanosome (brown).

In total, Orf3 affected 71 DS-affinity proteins identified from A549 cells, which includes those directly interacting with Orf3 and those perturbed by Orf3 protein expression in A549 cells. The large number of Orf3-affected host proteins implicates important roles of Orf3 in SARS-CoV-2 infection. Network analysis reveals these proteins to be mostly associated with gene expression regulation, cytoplasmic vesicles, apoptosis, response to stress, monosaccharide biosynthesis, or hydrolase activity (Fig. 5). Several of these are classical nuclear autoAgs, e.g., PNCA, SSB (Lupus La), XRCC5 (Lupus Ku80, thyroid-lupus autoAg), XRCC6 (Ku70), and SNRPB (SmB/B’). A few are unknown autoAgs but with important relevance to COVID, e.g., PAFAH1B2 and PAFAH1B3 (the alpha catalytic subunits of the cytosolic type I platelet-activating factor (PAF) acetylhydrolase). PAF is produced by a variety of cells involved in host defense, and PAF signaling can trigger inflammatory and thrombotic cascades. The modulation of PAF by SARS-CoV-2 Orf3 may partially explain the frequently occurring thrombotic complications and coagulopathy in COVID-19 patients. PAF also induces apoptosis in a PAF receptor independent pathway that can be inhibited by PAFAH1B2 and PAFAH1B3 (38).

SARS-CoV-2 E protein affects a number of ribonucleoproteins that are related to translation initiation and mRNA splicing, e.g., hnRNP (U and UL1) and ribosomal protein (L7, L8 L11, L12, L35A). E-affected proteins are associated with establishment of protein localization to membrane, regulation of autophagy, and post-translational protein modification (Fig. 5). SARS-CoV-2 M, Nsp1, and N proteins also affect various ribonucleoproteins, whereas Nsp13 appears to affect proteins associated with the cytoskeleton. Overall, the majority of DS-affinity proteins found affected by individual SARS-CoV-2 proteins are known autoAgs (Fig. 5 and Table 1), which indicates that host proteins perturbed by viral components are an important source of autoAgs.

### Mitochondrial perturbation by SARS-CoV-2 Orf9b

By DS-affinity fractionation, we identified 22 mitochondrial ribosomal proteins from A549 cells with strong DS-affinity, including mitochondrial 39S ribosomal proteins (L1, L2, L13, L15, L17, L18, L19, L23, L37, L38, L39, L45, L49) and 28S ribosomal proteins (S9, S22, S23, S27, S28, S29, S30, S34, S39) (Table 1). Remarkably, 14 mitochondrial ribosomal proteins are found down-regulated in SARS-CoV-2 infection, with 2 (L15 and L37) reported both down- and up-regulated. Moreover, MRPS27 is found in the interactome of SAS-Cov-2 Nsp8. Most strikingly, Orf9b-expression in A549 cells caused down-regulation of 16 proteins that we identified by DS-affinity, namely, ALB, ANXA4, LAMP2, MRPL2, MRPL13, MRPL15, MRPL17, MRPL18, MRPL19, MRPS30, MRPL37, MRPL38, MRPL39, MRPL45, MRPL49, and UCHL1 (Fig. 5). Eleven of the Orf9-affected proteins are mitochondrial ribosomal proteins, which may affect the mitochondrial translation machinery. Orf9-affected proteins may also be involved in mitochondrion localization, autophagosome maturation, or other processes. Overall, these findings suggest that SARS-CoV-2 infection may affect mitochondria primarily through Orf9b.

Orf9b of SARS-CoV has been shown to localize to mitochondria, trigger ubiquitination and proteasomal degradation of dynamin-like protein 1, limit host cell interferon signaling by targeting mitochondrial associated adaptor molecule MAVS signalosome, and manipulate the mitochondrial function to help evade host innate immunity (39). Orf9b of SARS-CoV-2 has been reported to suppress the type I interferon response by targeting TOM70 (40). In COVID-19 pneumonia patients, monocytes show altered bioenergetics and mitochondrial dysfunction with depolarized and abnormal ultrastructure (41).

Currently, little is known about the involvement of mitochondrial ribosomes or mitochondrial translation in SARS-CoV-2 infection. Expression of mitochondrial ribosomal proteins associated with protein synthesis has been found to be the most striking transcriptional difference among dengue virus-infected children, as revealed by a genome-wide microarray analysis of whole blood RNA from 34 infected children collected on days 3-6 of illness (42). In human cytomegalovirus infection, proteins involved in biogenesis of the mitochondrial ribosome changed early during the viral replication cycle (43). Mitochondria are vital to cell survival and apoptosis as they produce the majority of adenosine triphosphate (ATP) that provide chemical energy to cells. Especially for cells such as muscles that require much ATP, mitochondrial dysfunction will certainly lead to problems such as muscle weakness and fatigue. The roles of mitochondrial ribosomal proteins play in COVID and long-term sequelae merit further investigation.

### AutoAgs related to ubiquitination alteration in SARS-CoV-2 infection

Ubiquitination provides a universal signal for protein degradation. By comparing our data with the ubiquitinome of SARS-CoV-2 infected cells, we identified 102 DS-affinity proteins that are altered by ubiquitination during viral infection (Supplemental Table 1). These ubiquitination-altered proteins are significantly associated with gene expression, catabolic process, regulation of apoptotic process, cytoplasmic vesicles, and cytoskeleton (Fig. 6). They include 15 ribosomal proteins, 8 heat shock proteins, 5 hnRNP proteins, 5 histones, 4 translation elongation factors, and 3 translation initiation factors, and a majority of them are known autoAgs (Table 1).

**Fig. 6.**
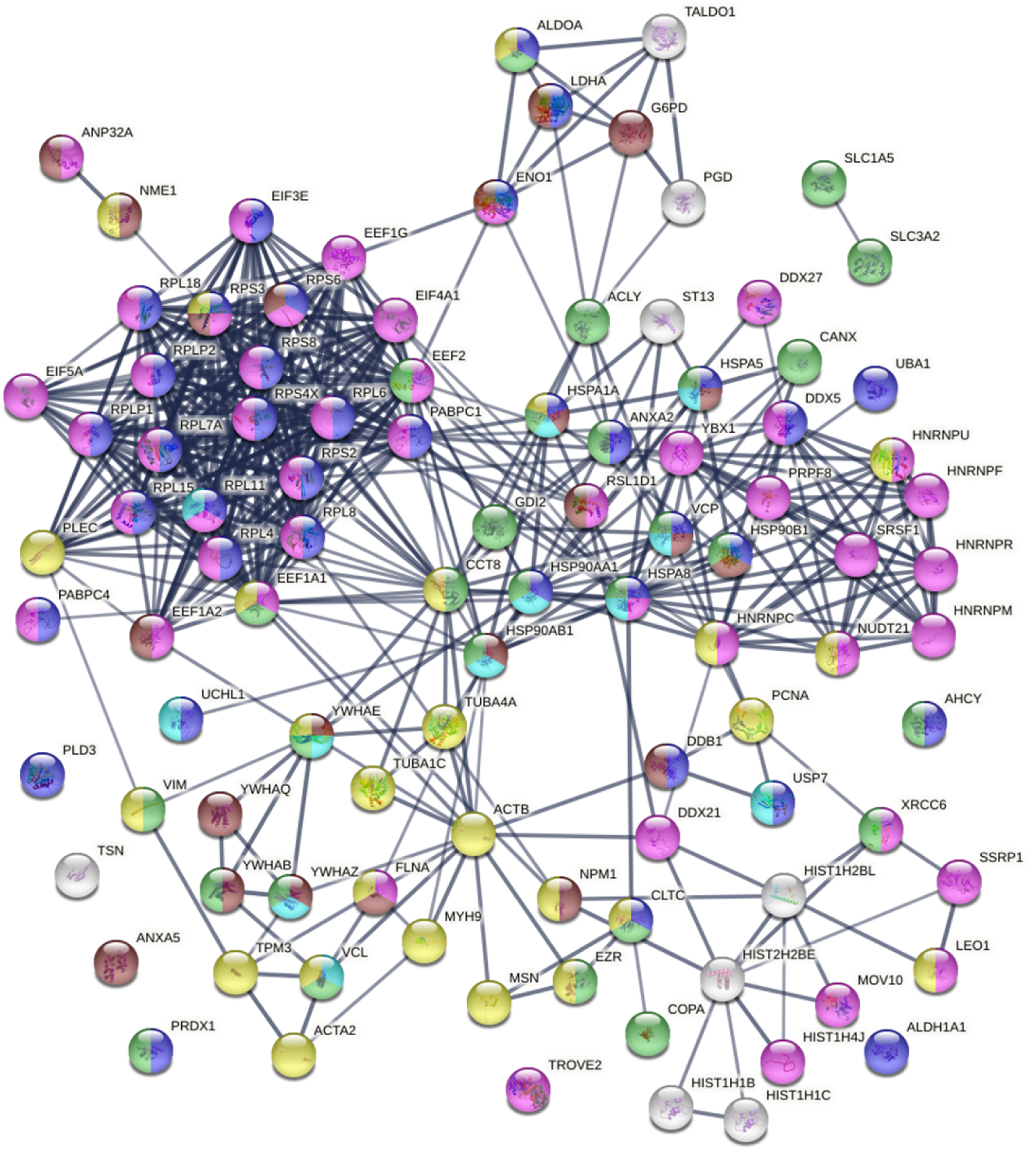
Known and putative autoAgs derived from ubiquitination alteration in SARS-CoV-2 infected AF549 cells. Marked proteins are associated with catabolic process (37 proteins, blue), ubiquitin protein ligase binding (12 proteins, aqua), gene expression (46 proteins, pink), regulation of apoptotic process (22 proteins, brown), cytoplasmic vesicles (28 proteins, green), and cytoskeleton (26 proteins, gold).

Three ubiquitination/de-ubiquitination enzymes (UBA1, UCHL1, and USP7) are COVID-altered and possess DS-affinity, with UBA1 and UCHL1 being known autoAgs. UBA1 catalyzes the first step in ubiquitin conjugation to mark proteins for degradation through the ubiquitin-proteasome system. USP7 is a hydrolase that deubiquitinates target proteins. UCHL1 is a thiol protease that recognizes and hydrolyzes a peptide bond at the C-terminal glycine of ubiquitin, and is involved in the processing of ubiquitin precursors and of ubiquitinated proteins. UBA1 is found down-regulated by Orf3 expression. UCHL1 is found in the Orf3 interactome, up-regulated by SARS-CoV-2 E protein expression, and down-regulated by Nsp12, Nsp8, Orf8, and Orf9b (Supplemental Table 1).

Ten COVID-altered DS-affinity proteins were identified with ubiquitin protein ligase-binding activity, including HSPA1A, HSPA5, HSPA8, HSP90AA1, HSP90AB1, RPL11, VCP, VCL, YWHAE, and YWHAZ. HSPA1A interacts with SARS-CoV-2 Orf9b and N proteins, HSPA5 (GRP78/BiP) interacts with Nsp2 and Nsp4, and HSPA8 interacts with Nsp2. Except for RPL11, all are known autoAgs (Table 1).

In addition, SARS-CoV-2 and other coronaviruses encode for papain-like proteases (PLP), an important multifunctional enzyme with de-ubiquitination, de-ISGlation, and interferon antagonism activities (44). PLPs, along with other proteases, are responsible for processing replicase proteins that are required from viral replication. PLP of SARS-CoV-2 is able to reverse host ubiquitination and remove interferon-stimulated gene product 15 (ISG15), and its substrate activity mirrors closely that of PLP of MERS (45). Ubiquitin modifications can regulate innate immune response and apoptosis, and ISG15 is a ubiquitin-like modifier typically expressed during host cell immune response. Overall, various components of SARS-CoV- 2 appear to be able to alter uniquitination of host proteins. The large pool of ubiquitin-altered proteins in SARS-CoV-2 infection indicates that ubiquitin modification, such as differential abundance and dynamic ubiquitination pattern change, may be a major origin of autoAgs.

### AutoAgs related to phosphorylation alteration in SARS-CoV-2 infection

Comparing our data with currently available phosphoproteome data of COVID-19 (26, 34), 97 phosphoproteins are found with both DS-affinity and COVID-induced alteration, with a majority (52/97) related to gene expression (Fig. 7). Notably, they include 8 heterogeneous nuclear ribonucleoproteins affected by phosphorylation (HNRNPA2B1, HNRNPC, HNRNPH1, HNRNPK, HNRNPM, HNRNPU, HNRNPUL1, HNRNPUL2), with 4 being known autoAgs (Table 1). HNRNPs are involved in many cellular processes, including gene transcription, cell cycle, DNA damage control, post-transcriptional modification of newly synthesized pre-mRNA, and virus replication. For example, HNRNPA2B1 displays RNA-binding affinity to murine hepatitis coronavirus via binding the viral RNA at the 3’ end and modulate viral RNA synthesis (46).

**Fig. 7.**
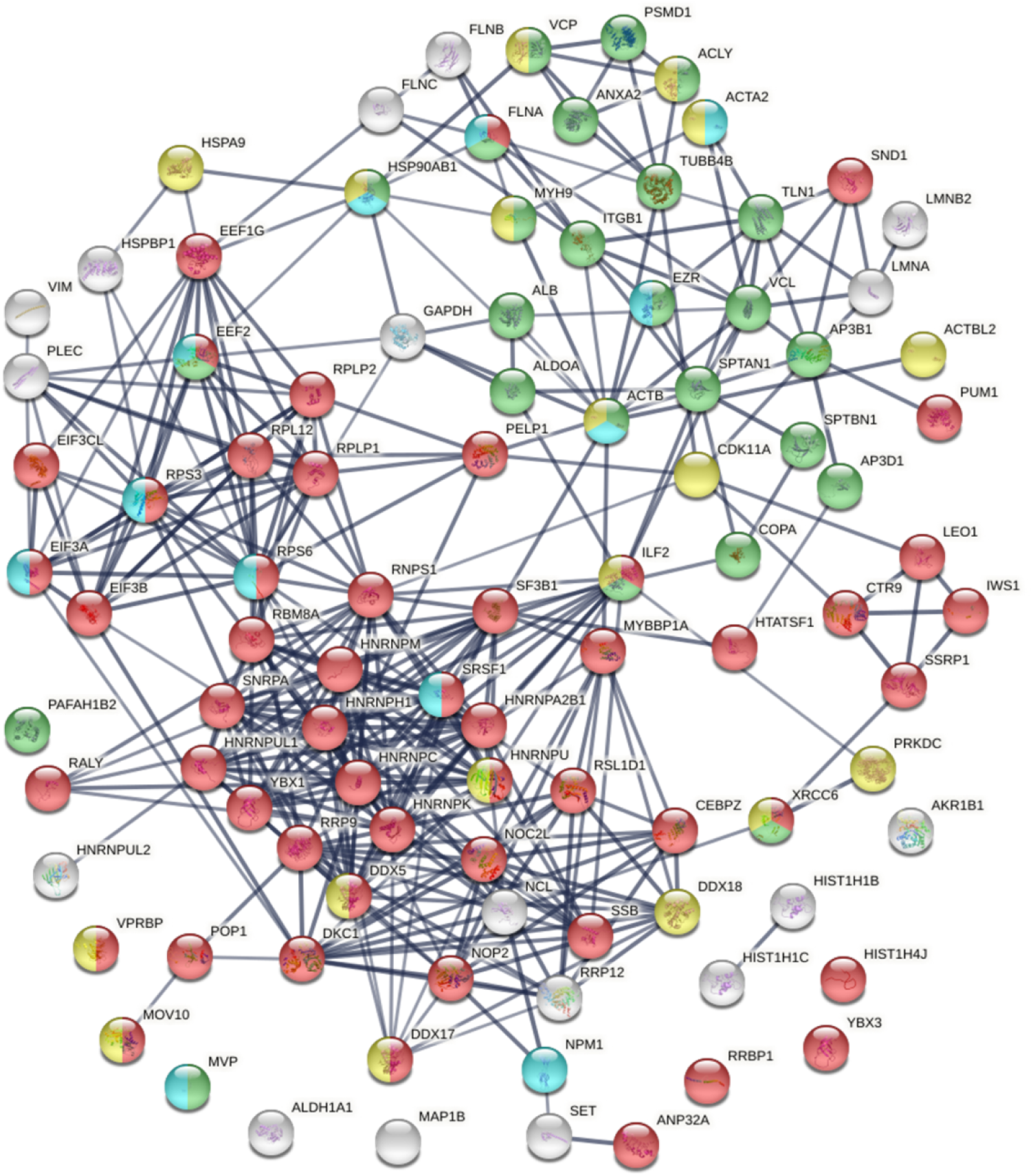
Known and putative autoAgs derived from phosphorylation alteration in SARS-CoV-2 infected cells. Marked proteins are associated with gene expression (52 proteins, red), vesicle-mediated transport (25 proteins, green), ATP binding (18 proteins, gold), and kinase binding (12 proteins, aqua).

There are 25 phosphorylation-altered proteins that are related to vesicle-mediated transport, most of which are known autoAgs, including ACLY, ACTA2, ACTB, ALB, ALDO, ANXA2, FLNA, COPA, SPTAN1, SPTBN1, TLN1, TUBB4, and VCL (Fig. 7 and Table 1). The coatomer, to which COPA (coatomer subunit alpha) belongs, is a cytosolic protein complex that associates with Golgi non-clathrin-coated vesicles and is required for budding from the Golgi membrane. COPA is associated with autoimmune interstitial lung, joint, and kidney disease (47).

There are 18 phosphorylation-altered potential autoAgs with ATP binding activity, and 12 with kinase binding activity (Fig. 7). In particular, PRKDC (DNA-dependent protein kinase catalytic subunit) is identified with strong DS-affinity and is a known autoAg. It is a serine/threonine-protein kinase that acts as a molecular sensor for DNA damage, with involvement in numerous biological processes such as DNA damage and repair, immunity, innate immunity, ribosome biogenesis, and apoptosis. PRDKC is found in the interactomes of M and Nsp4 proteins of SARS-CoV-2 and up-regulated by expression of Nsp10, Nsp9, Orf7a, or Orf7b protein in A549 cells (19, 34). PRKDC is also found up-regulated at 0 h and 4 h in SARS-CoV-2 infected Vero E6 cells (26) and up-regulated at 24 h in SARS-CoV-2 infected Caco-2 cells (20). These findings suggest that phosphorylation by PRKDC plays extensive and important roles in COVID.

Proteins phosphorylated during apoptosis are common targets of autoantibodies. For example, the U1- 70 snRNP autoAg undergoes specific changes in the phosphorylation/dephosphorylation balance and cellular localization during apoptosis (48), and phosphorylated U1-snRNP complex induced by apoptosis is recognized by autoantibodies in patients with systemic lupus erythematosus (49). A high degree of phosphorylation of SSB (lupus La autoAg) substantially diminished its poly(U) binding capacity, but its binding to human autoantibodies increased 2-fold with increased phosphorylation (50). On the other hand, SSB autoAg has also been reported to be dephosphorylated and cleaved during early apoptosis (51). During apoptosis, ribosomal protein P1 and P2 autoAgs are completely dephosphorylated while P0 autoAg is partially dephosphorylated (52). Therefore, alterations in phosphorylation, either hyper- or hypo- phosphorylation, may lead to changes in self-molecules and render them autoantigenic.

## Conclusion

In our quest for a comprehensive autoantigen atlas for COVID-19, we report an autoantigen profile of 191 confirmed autoAgs and 150 putative autoAgs in SARS-CoV-2 infection. These proteins are initially identified from human lung epithelial A549 cells using a unique DS-affinity autoAg enrichment strategy, and then compared with currently available COVID-omics data. Our study reveals that cellular processes and components integral to viral infection are major origins of autoAgs, including gene expression, ribonucleoprotein biogenesis, translation and mitochondrial translation, vesicle and vesicle-mediated transport, and cytoskeleton. Ubiquitination and phosphorylation are particular post-translational modifications that cause changes in self-molecules and render them autoantigenic. Impaired clearance of apoptotic and dead cell material is considered a major pathogenic attribute to autoimmune disease. We have previously shown that DS possesses unique affinity to apoptotic cells and their released autoAgs, and our current study further demonstrates that ubiquitination and phosphorylation associated with apoptosis are possibly major sources of molecular alterations in self-molecule to autoantigen transformation. Overall, our study demonstrates that SARS-CoV-2 causes extensive alterations of host cellular proteins and produces a large number of potential autoAgs, indicating that there may be an intimate relationship between COVID infection and autoimmunity.

## Materials and Methods

### A549 cell culture

The A549 cell line was obtained from the ATCC (Manassas, VA, USA) and cultured in complete F-12K medium at 37 °C in 75 cm^2^ flasks to 80% confluency. The growth medium was supplemented with 10% fetal bovine serum and a penicillin-streptomycin-glutamine mixture (Thermo Fisher).

### Protein extraction

About 100 million A549 cells were suspended in 10 ml of 50 mM phosphate buffer (pH 7.4) containing the Roche Complete Mini protease inhibitor cocktail. Cells were homogenized on ice with a microprobe sonicator until the turbid mixture turned nearly clear with no visible cells left. The homogenate was centrifuged at 10,000 g at 4 °C for 20 min, and the total protein extract in the supernatant was collected. Protein concentration was measured by absorbance at 280 nm using a NanoDrop UV-Vis spectrometer (ThermoFisher).

### DS-Sepharose resin preparation

The DS-affinity resins were prepared as previously described (3, 5). In brief, 2 ml of EAH Sepharose 4B resins (GE Healthcare Life Sciences) were washed with distilled water three times and mixed with 10 mg of DS (Sigma-Aldrich) in 1 ml of 0.1 M MES buffer, pH 5.0. About 20 mg of N-(3-dimethylaminopropyl)-N’- ethylcarbodiimide hydrochloride (Sigma-Aldrich) powder was added at the beginning of the reaction, and another 20 mg was added after 8 h of reaction. The reaction mixture was mixed by end-over-end rotation at 25 °C for 16 h. The coupled resins were washed with water three times and equilibrated with a low-pH buffer (0.1 M acetate, 0.5 M NaCl, pH 5.0) and a high-pH buffer (0.1 M Tris, 0.5 M NaCl, pH 8.0).

### DS-affinity fractionation

The total proteins extracted from A549 cells were fractionated in a DS-Sepharose column with a BioLogic Duo-Flow system (Bio-Rad). About 40 mg of proteins in 40 ml of 10 mM phosphate buffer (pH 7.4; buffer A) were loaded onto the column at a rate of 1 ml/min. Unbound and weakly proteins were washed off with 60 ml of buffer A and then 40 ml of 0.2 M NaCl in buffer A. The remaining bound proteins were eluted with 40 ml 0.5 M NaCl and then with 40 ml 1.0 M NaCl in buffer A. Fractions were desalted and concentrated to 0.5 ml with 5-kDa cut-off Vivaspin centrifugal filters (Sartorius). Fractionated proteins were separated by 1-D SDS-PAGE in 4-12% Bis-Tris gels, and the gel lanes were divided into two or three sections and subjected to sequencing.

### Mass spectrometry sequencing

Protein sequencing was performed at the Taplin Biological Mass Spectrometry Facility at Harvard Medical School. Proteins in gels were digested with sequencing-grade trypsin (Promega) at 4 °C for 45 min. Tryptic peptides were separated on a nano-scale C_18_ HPLC capillary column and analyzed in an LTQ linear ion-trap mass spectrometer (Thermo Fisher). Peptide sequences and protein identities were assigned by matching the measured fragmentation pattern with proteins or translated nucleotide databases using Sequest. All data were manually inspected. Only proteins with ≥2 peptide matches were considered positively identified.

### COVID data comparison

DS-affinity proteins were compared with currently available proteomic and transcriptomic data from SARS-CoV-2 infection compiled in the Coronascape database (as of 12/14/2020) (17-37). These data had been obtained with proteomics, phosphoproteomics, interactome, ubiquitome, and RNA-seq techniques. Up- and down-regulated proteins or genes were identified by comparing COVID-19 patients vs. healthy controls and cells infected vs. uninfected by SARS-CoV-2. Similarity searches were conducted between our data and the Coronascape database to identify DS-affinity proteins (or their corresponding genes) that are up- and/or down-regulated in the viral infection.

### Protein-protein interaction network analysis

Protein-protein interactions were analyzed by STRING (16). Interactions include both direct physical interaction and indirect functional associations, which are derived from genomic context predictions, high-throughput lab experiments, co-expression, automated text mining, and previous knowledge in databases. Each interaction is annotated with a confidence score from 0 to 1, with 1 being the highest, indicating the likelihood of an interaction to be true. Only interactions with high confidence (a minimum score of 0.7) are shown in the figures.

### Pathway and process enrichment analysis

Pathways and processes enrichment were analyzed with Metascape (17), which utilize various ontology sources such as KEGG Pathway, GO Biological Process, Reactome Gene Sets, Canonical Pathways, CORUM, TRRUST, and DiGenBase. All genes in the genome were used as the enrichment background. Terms with a p-value <0.01, a minimum count of 3, and an enrichment factor (ratio between the observed counts and the counts expected by chance) >1.5 were collected and grouped into clusters based on their membership similarities. The most statistically significant term within a cluster was chosen to represent the cluster. Hierarchical clustering trees were obtained with ShinyGo (53).

### Autoantigen confirmation literature text mining

Literature searches in Pubmed were performed for every DS-affinity protein identified in this study. Search keywords included the protein name, its gene symbol, alternative names and symbols, and the MeSH keyword “autoantibodies”. Only proteins with their specific autoantibodies reported in PubMed-listed journal articles were considered “confirmed” autoAgs in this study.

## Acknowledgements and funding statement

This work was partially supported by Curandis, the US NIH, and a Cycle for Survival Innovation Grant (to MHR). MHR acknowledges the NIH/NCI R21 CA251992 and MSKCC Cancer Center Support Grant P30 CA008748. The funding bodies were not involved in the design of the study and the collection, analysis, and interpretation of data. We thank Jung-hyun Rho for technical assistance with experiments. We thank Ross Tomaino and the Taplin Biological Mass Spectrometry facility of Harvard Medical School for expert service with protein sequencing.

## Authors’ contributions

JYW directed the study, analyzed data, and wrote the manuscript. WZ performed some experiments and reviewed the manuscript. MWR and VBR assisted with data analysis and manuscript preparation. MHR consulted on the study and data analysis and edited the manuscript. All authors have approved the manuscript.

